# Nutrient-driven dedifferentiation of enteroendocrine cells promotes adaptive intestinal growth

**DOI:** 10.1101/2023.05.08.539820

**Authors:** Hiroki Nagai, Luis Augusto Eijy Nagai, Sohei Tasaki, Ryuichiro Nakato, Daiki Umetsu, Erina Kuranaga, Masayuki Miura, Yu-ichiro Nakajima

## Abstract

Post-developmental organ resizing improves organismal fitness under constantly changing nutrient environments. Although stem cell abundance is a fundamental determinant of adaptive resizing, our understanding of its underlying mechanisms remains primarily limited to the regulation of stem cell division. Here we demonstrate that nutrient fluctuation induces dedifferentiation in the *Drosophila* adult midgut to drive adaptive intestinal growth. From lineage tracing and single-cell RNA-sequencing, we identify a subpopulation of enteroendocrine cells (EEs) that convert into functional intestinal stem cells (ISCs) in response to dietary glucose and amino acids by activating the JAK-STAT pathway. Genetic ablation of EE-derived ISCs severely impairs ISC expansion and midgut growth despite the retention of resident ISCs, and *in silico* modeling further indicates that EE dedifferentiation enables efficient increase in the midgut cell number while maintaining epithelial cell composition. Our findings uncover a physiologically-induced dedifferentiation that ensures ISC expansion during adaptive organ growth in concert with nutrient conditions.

## INTRODUCTION

Adult organs in Metazoa flexibly remodel their structure in response to environmental factors. In particular, the intestine adapts to nutrient availability by dynamically changing its organ size: the intestine shrinks during starvation and enlarges upon refeeding, which optimizes digestive and absorptive performance^1–6^. Such adaptive resizing is crucial for organ fitness and health since failure in regrowth leads to pathologies such as short bowel syndrome^6, 7^. It should be noted that most adult organs harbor regional differences in cellular composition and functions^8–13^, implying that the mechanisms driving the adaptive responses are diversified across distinct regions. Although the abundance of stem cells is a fundamental determinant of organ size^3, 14, 15^, it remains largely unknown how the organ-wide expansion of the stem cell pool is coordinated in different regions and achieved during adaptive resizing.

Accumulating evidence has revealed that daughters of tissue stem cells exert differentiation plasticity under severely stressful conditions: the stem cell pool can be restored even after their complete loss through the reversion of differentiated cells into functional stem cells. This cell fate plasticity, hereafter called dedifferentiation, was initially identified upon lens removal in newt, and is now recognized as an evolutionary conserved regenerative strategy that revives lost stem cells^14, 16–18^. In mammals, dedifferentiation has been identified in multiple tissues, among which the intestinal epithelium exhibits a highly plastic nature: both absorptive and secretory lineages undergo dedifferentiation into intestinal stem cells (ISCs) upon severe injury or during inflammatory tumorigenesis^19–26^. However, current observations of cell fate plasticity have been limited to experimental systems either wherein near-total active stem cells are eliminated or in pathological contexts^16, 18^. It thus remains largely unclear whether cell fates are plastic under physiological conditions or as the result of naturally occurring perturbations.

The cellular lineage of the adult intestinal epithelium is highly conserved between *Drosophila* and mammals^27–29^. In the *Drosophila* adult midgut, asymmetric division of an ISC generates another ISC and a progenitor, either an enteroblast (EB) or an enteroendocrine progenitor (EEP); then the EB or the EEP differentiates into an absorptive enterocyte (EC) or a secretory EE, respectively. After the eclosion of adult flies, the number of ISCs, as well as the total cell number, dramatically increases in a feeding-dependent manner (Figures 1A, 1B and S1A-S1F), driving adaptive intestinal growth^3^. Previous reports have shown that food intake induces symmetric ISC division via insulin signaling in the posterior region of the midgut^3, 30–32^, but whether self-renewal of ISCs is the sole mechanism for ISC expansion in the rapidly growing midgut remains unclear.

**Figure 1.**
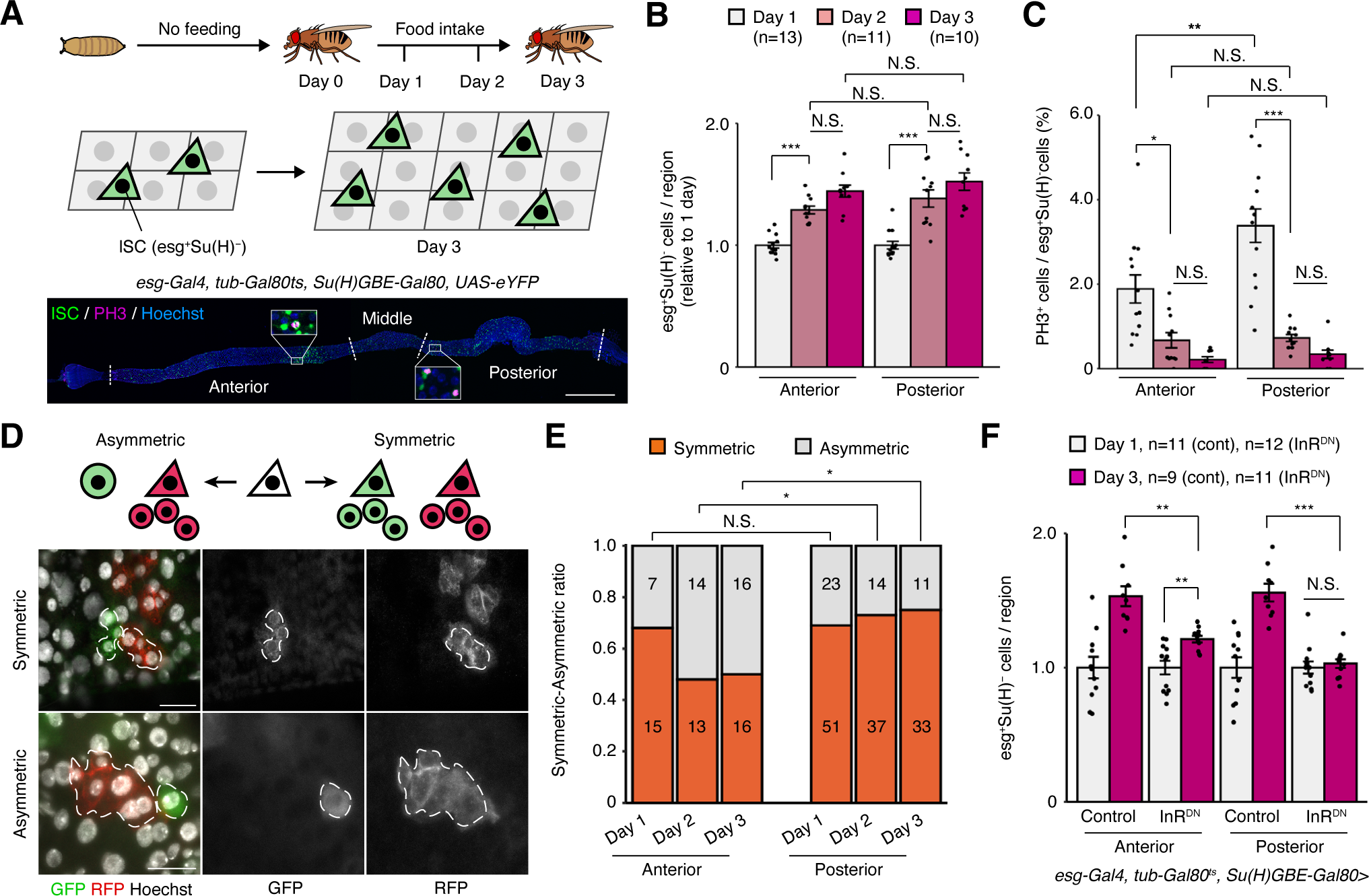
Self-renewal of ISCs is not sufficient for ISC expansion in the anterior midgut. (A) Schematic of ISC expansion in early adult stage. The anterior, middle, and posterior region of the adult midgut is indicated in the confocal image (see also STAR Methods for determination of regional boundaries). (B) The relative increase of *esg^+^Su(H) ^−^* cell number between Day 1 and Day 3 guts. (C) The mitotic activity of *esg^+^Su(H)^−^* cells. The same samples are quantified in (B) and (C). (D) Twin spot MARCM technique labels one ISC daughter with GFP and the other with RFP. In the case of symmetric division, both ISC daughters generate additional cells, resulting in multiple cells both in the GFP and the RFP clones. In the case of asymmetric division, one daughter differentiates and stops mitosis while the other daughter continues to proliferate, resulting in one clone with one cell and the other with multiple cells. Please see also STAR Methods for the classification of symmetric/asymmetric division. (E) The ratio of symmetric/asymmetric ISC division in the Day 1, 2, and Day 3 midgut. (F) The relative increase of *esg^+^Su(H)^−^*cell number in midguts overexpressing the dominant negative form of InR (InR^DN^). N.S., not significant: P>0.05, *P≤0.05, **P≤0.01, ***P≤0.001. One-way ANOVAs with post hoc Tukey test (B, C, F), chi-square test (E). *n* indicates the number of midguts in (B, C, F) and the number of clones in (E). Scale bars: 500 µm (A), 20 µm (D). See also Figure S1.

Here, we investigate the potential involvement of cell fate plasticity in nutrient-dependent ISC expansion and subsequent intestinal growth using the *Drosophila* adult midgut. In contrast to the posterior midgut where symmetric ISC division fuels stem cell pool replenishing, we show that a subset of EEs frequently dedifferentiate into functional ISCs in response to nutritional stimuli in the anterior midgut. Single-cell transcriptome and *in vivo* lineage tracing identify AstC (somatostatin in mammals) positive EEs as the EE subpopulation exhibiting high cell fate plasticity in the early adult midgut. We further reveal that EE dedifferentiation functions as an irreplaceable source of additional ISCs and thus drives intestinal growth. Notably, a starvation-refeeding cycle also induces the EE-to-ISC conversion in mature adults, indicating that EE dedifferentiation generally occurs in response to nutrient fluctuation. These results demonstrate the nutritional regulation of and the role of dedifferentiation in physiologically induced stem cell expansion.

## RESULTS

### Self-renewal of ISCs is not sufficient for nutrient-dependent ISC expansion in the anterior midgut

To test whether stem cell expansion is entirely dependent on symmetric ISC division, we first examined the mitotic activity of ISCs. To this end, we used a known ISC marker, *esg^+^Su(H)^−^*, and counted the number of *esg^+^Su(H)^−^* cells as well as the number of mitotic marker (phosphohistone H3; PH3) positive cells in whole mount midguts by labeling *esg^+^Su(H)^−^*cells with the GAL4/UAS system (*esg-Gal4, tub-Gal80^ts^, Su(H)GBE- Gal80>UAS-eYFP*) (Figure 1A). The number of *esg^+^Su(H)^−^* cells increased by ∼1.5 fold in both anterior and posterior regions during the first three days of the adult stage (Figures 1B and S1B). However, the PH3^+^ ratio of *esg^+^Su(H)^−^* cells in the anterior midgut was significantly lower than that of the posterior midgut at 1-day-old (Day 1, Figure 1C), suggesting distinct mitotic activity between anterior and posterior ISCs. We confirmed these results utilizing the Gal4 driver of another ISC marker, *Dl* (Figure S1C), and using the endogenously GFP-tagged protein trap line *esg-GFP* (Figures S1D-S1F).

Despite lower mitotic activity in the anterior midgut, the increase in ISC number is comparable between the two regions (Figure 1B). One explanation for this finding is that anterior ISCs more preferentially divide symmetrically than do posterior ISCs in order to increase their number. To test this possibility, we generated twin-spot clones using the mosaic analysis with a repressible cell marker (MARCM) technique that allows for the identification of asymmetric and symmetric cell division of ISCs^33^ (Figures 1D and S1H). The proportion of symmetric ISC division in the anterior region was comparable to or even lower than that in the posterior region throughout the first three days after eclosion (Figure 1E), suggesting the existence of other mechanisms that contribute to ISC expansion in the anterior midgut beyond symmetric division. Consistent with this observation, induction of the dominant negative form of the insulin receptor, which strongly blocks nutrient-dependent ISC division^34, 35^, only partially suppressed stem cell expansion in the anterior region, while almost completely eliminating ISC expansion in the posterior region (Figure 1F). These results suggest that symmetric ISC division alone does not account for ISC expansion in the anterior midgut, raising the possibility of as-yet unidentified cell fate reversion during nutrient-dependent intestinal growth.

### Apoptosis-independent decline in EE number during midgut growth

While the number of EBs, the enterocyte progenitor, increased both in the anterior and the posterior region in the early adult intestine^3^ (Figure S1G), the dynamics of EEs are unclear. We thus decided to explore the number of EEs under two conditions: using the EE-specific driver *pros-Gal4* (*pros-Gal4>UAS-GFP*) and with anti-Pros staining for the wild-type fly. We found that EE population significantly decreased during the first three days of adult life, and then recovered on Day 7 (Figures 2A and S2A). Notably, the decline in EE number was a feeding-dependent process, and was more prominent in the anterior midgut, where we have established that self-renewal of ISCs is insufficient for the expansion of the stem cell pool (Figures 2A, 2B, S2A, and S2B). We then tested the possibility that EEs undergo apoptosis, but found that EEs rarely exhibited cell death markers (Figures S2C-S2F). Furthermore, overexpression of apoptosis inhibitors *p35* or *diap1* by *pros-Gal4* failed to suppress the decline of EE number (Figures S2G and S2H). Together, these results excluded apoptosis as the cause of the EE decrease and led us to hypothesize that cell fate conversion from EE to ISC underlies ISC expansion.

**Figure 2.**
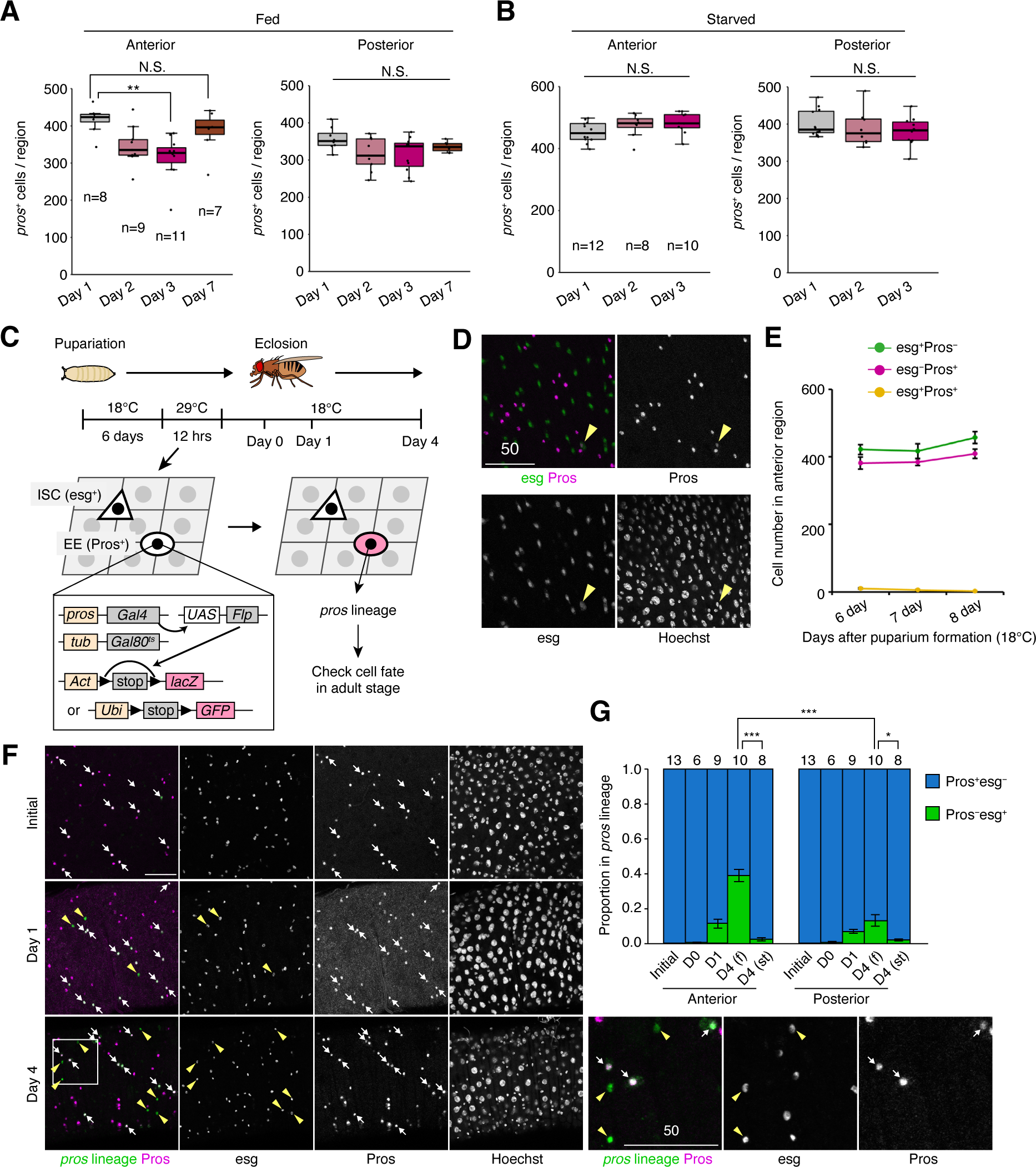
A subset of EEs dedifferentiates into ISCs in response to food intake after eclosion. (A and B) The number of *pros-Gal4>UAS-GFP^+^*EEs in Day 1, 2, 3 and Day 7 fed (A) and starved (B) guts. (C) Schematic of lineage tracing. Adult EEs were labeled with lacZ (β-gal) or GFP before eclosion, and their cell fate was checked after eclosion. (D) Representative image of Pros^+^ cells and esg-GFP^+^ cells in the midgut at 6 days after puparium formation. Arrowhead indicates Pros^+^esg^+^ cell. (E) Quantification of Pros^−^esg^+^ cells (green), Pros^+^esg^−^ cells (magenta), and Pros^+^esg^+^ cells (yellow) at 6, 7, and 8 days after puparium formation. (F) Representative images of lineage tracing. Arrows: *pros-*lineage^+^Pros^+^esg^−^ cells, arrowheads: *pros-*lineage^+^Pros^−^esg^+^ cells. (G) Quantification of Pros^+^esg^−^ and Pros^−^esg^+^ ratio in *pros*-lineage cells. Both fed (f) and starved (st) conditions were assessed for Day 4. N.S., not significant: P>0.05, *P≤0.05, **P≤0.01, ***P≤0.001. One-way ANOVAs with post hoc Tukey tests. *n* indicates the number of midguts. Scale bars: 50 µm. See also Figure S2 and S3.

### A subset of EE converts into functional ISCs in response to food intake

To investigate cell fate dynamics of EEs after eclosion, we performed a lineage tracing experiment in which temperature shift induces permanent labeling of *pros^+^* EE-derived cells with GFP or lacZ (Figure 2C)^36, 37^. Since the formation of adult differentiated EEs (Pros^+^esg^−^, Pros^+^piezo^−^, or Pros^+^Dl^−^ cells) is completed during the pupal stage (Figures 2D, 2E and S3A-S3D)^38–40^, we labeled EEs before eclosion and examined their cell fate in the adult stage by examining expression of Pros and the stemness marker *escargot* (*esg*)^41^ (Figure 2C). We first confirmed that our scheme specifically labeled Pros^+^esg^−^ cells at the beginning of lineage tracing (Figure 2F, 2G, and S3E; 100% of labeled cells were Pros^+^esg^−^ in 11/13 midguts). While 99.3 ± 0.3% of traced cells maintained Pros expression just after eclosion (Day 0), 9.7 ± 1.8% of *pros*-lineage cells lost Pros signal and acquired expression of esg in Day 1 adults, and this proportion reached 27.3 ± 3.0% in Day 4 adults (Figure 2F and 2G). The lineage-traced Pros negative cells also expressed another ISC marker, *Delta (Dl)*, but rarely expressed the EB marker *Su(H)* (Figure S3F-S3H), suggesting the direct conversion of EEs into a stem-like state. Importantly, induction of the *pros*-derived *esg^+^* population was dependent on food intake and was more frequent in the anterior region (Figure 2G), similar to the dramatic decline in EE number in the anterior midgut (Figures 2A, 2B, S2A, S2B). These results indicate that the first food intake after eclosion induces cell fate reversion from EE to ISC.

To examine how EEs lose their identity and acquire ISC fate, we first monitored the dedifferentiation process after feeding. In the young adult midgut, typical cellular morphology delimited by anti-Armadillo staining is round for EEs and angular for ISCs/EBs (Figure 3A)^42–44^. Interestingly, we found that *pros*-lineage cells transform their morphology after acquiring *esg* expression: while the *pros*-derived *esg^+^* cells exhibited a rounded shape in Day 1 guts, their shape became angular in Day 4 guts (Figures 3A, 3B, and S3I). We also found that remnants of neuropeptide CCHa1 persist in *pros*-lineage *esg^+^* cells in the Day 1 guts but disappear in the Day 4 guts (Figures 3C and S3J), suggesting that these *esg^+^* cells originated from mature EEs. These results together indicate that characteristics of EEs are gradually lost in the fate converting cells, which is consistent with the gradual transcriptional repression of dedifferentiating secretory cells in the regenerating mammalian intestine^22^.

**Figure 3:**
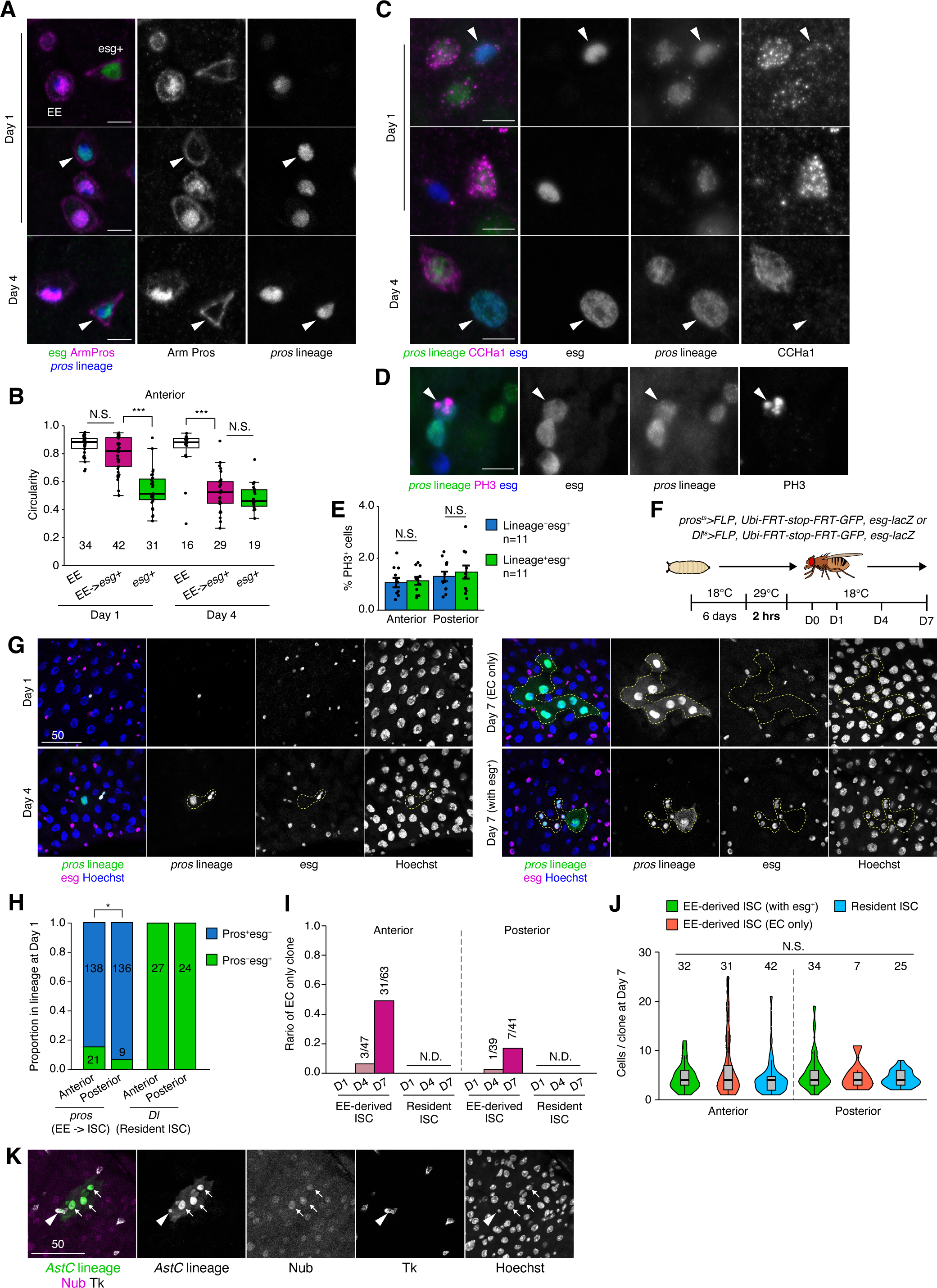
*pros*-derived Pros^−^esg^+^ cells are functional ISCs. (A) Histological analysis of cellular shape. Lineage tracing of EEs was performed, and the shape of EEs (Pros^+^esg^−^), EE-derived esg^+^ cells (*pros-*lineage^+^Pros^−^esg^+^, arrowheads), and non-EE-derived esg^+^ cells (*pros-*lineage ^−^Pros ^−^esg^+^) was examined by anti-Arm staining that visualizes adherens junction. (B) Quantification of (A). Circularity of EEs, EE-derived esg^+^ cells, and non-EE-derived esg^+^ cells in the anterior region were quantified. (C) EE-derived esg^+^ cells (arrowheads) contained the remnants of the CCHa1 peptide in Day 1 fed guts but not in Day 4 fed guts. (D) PH3 signal in EE-derived esg^+^ cells. (E) The mitotic activity of EE-derived esg^+^ cells was comparable to that of conventional (non-EE-derived) esg^+^ cells. PH3 staining was performed after paraquat feeding (5 mM, Day 4-5). (F) Schematic for sparse labeling. Two hours incubation at 29°C sparsely labeled EE lineage cells (*pros*) and resident ISC lineage cells (*Dl*). (G) Representative images of EE-derived esg^+^ cell clone at Day 1, 4, and 7. The clone containing only polyploid ECs (EC only) and the one retaining esg^+^ cells (with esg^+^) are shown for Day 7. (H) Quantification of Pros^+^esg^−^ and Pros^−^esg^+^ ratio in EE lineage and resident ISC lineage at Day 1. (I) The ratio of EC-only clones in lineage traced clones. (J) The number of cells per clone at Day 7 for each clone type. (K) Nub^+^ECs and Tk^+^EE in one clone that derived from AstC^+^EE. Arrows: Nub^+^ECs, arrowhead: Tk^+^EE. N.S., not significant: P>0.05, *P≤0.05, ***P≤0.001. Two tailed *t* tests (E), one-way ANOVAs with post hoc Tukey test (B, J), chi-square test (H). *n* indicates the number of cells (B), guts (E), and clones (H-J). Scale bars: 5 µm (A, C, D), 50 µm (G, K). See also Figure S3.

We next investigated whether the EE-derived stem-like cells exhibit proliferative capacity and generate differentiated daughter cells. We detected PH3 signal in EE-derived *esg^+^* cells with a frequency comparable to non-EE-lineage ISCs (resident ISCs, Figures 3D and 3E). To further examine the clonal expansion of EE-derived *esg^+^* cells and compare their behavior with resident ISCs, we sparsely labeled *pros-*lineage *esg^+^* cells as well as *Dl-*lineage cells before eclosion, and observed clones at several time points (Days 1, 4, 7; Figures 3F-3H). All *Dl*-lineage cells were Pros^−^esg^+^ at Day 1, confirming that they represented resident ISCs (Figure 3H). The number of cells per clone was comparable between the two stem cell populations (Figure S3K), but the clonal cell composition was distinct between them: a subset of EE-derived *esg^+^* cells, but none of the *Dl*-lineage resident ISCs, completely differentiated into *esg^−^* polyploid ECs at Day 7 (Figures 3G and 3I). Although the EC-only clones lost *esg^+^* cells, their cell number was similar to those retaining *esg^+^* cells (Figure 3J), suggesting that the EC-only clones were generated after several rounds of mitotic division. Moreover, the EE-derived clones that retained *esg^+^* cells also exhibited a higher ratio of ECs at the expense of *esg^+^* cells (Figures S3L and S3M). These results suggest that the EE-derived *esg^+^* cells have a differentiation bias toward ECs compared to resident ISCs. Notably, the ratio of the EC-only clones was considerably higher in the anterior midgut than the posterior midgut (Figure 3I), indicating the regional differences in the regulation of stem cell fate.

While a subset of EE-derived clones eventually became exclusively ECs, we also observed clones containing *esg^−^* diploid cells that are likely EEs (Figure S3N). To test the multipotency in the EE-derived *esg^+^*cells directly, we traced AstC^+^EE lineage and assessed the EC marker Nubbin as well as the EE marker Tk. Nubbin^+^ECs were detected in *AstC*-derived multicellular clones (Figure 3K), and EC character was further confirmed using the Myo31DF (Myo1A) reporter (Figure S3O)^45^. Furthermore, Tk^+^EE was also detected in the *AstC-*derived clones (Figure 3K). Given that the expression of AstC and Tk are mutually exclusive in differentiated EEs^46, 47^, the Tk^+^EE should be newly generated from AstC^+^EE-derived stem-like cells. Based on these observations, we concluded that the EE-derived *esg^+^* cells are multipotent ISCs that preferentially generate new ECs.

### Single-cell RNA sequencing identified a subpopulation of EEs undergoing dedifferentiation

To corroborate the dedifferentiation program of EEs with transcriptional profiling, we performed single-cell RNA sequencing (scRNA-seq) for the whole midgut samples from Day 1 and Day 3 young adults (Figures 4A and 4B). Transcriptome analysis of 4,184 high-quality cells (see STAR Methods) revealed 10 clusters that we annotated individually using known cell type-specific markers (Table S1) and validated by integrating with a published cell atlas from the Day 7 midgut^48^ (Figures S4A-S4C). Within the UMAP plot, ISCs and EEs in our scRNA-seq data formed two clusters each: ISC1 and ISC2 as well as AstC^+^EE and Tk^+^EE, respectively (Figures 4A-4C). ISC1 differentially expressed genes over ISC2 were enriched for GO terms related to cellular processes involved in the activation of tissue stem cells across species (Figure S4D)^49–53^. AstC^+^EE and Tk^+^EE are the major subclasses of EEs whose neuropeptide expression patterns are well recapitulated in our data (Figure S4C)^46, 48^. Notably, the ISC marker *Dl* was highly expressed in AstC^+^EEs (Figure 4C), and the AstC^+^EE gene signature was enriched for stem cell maintenance over Tk^+^EEs (Figures S4E-S4G). In addition, a portion of AstC^+^EEs, largely derived from the Day 1 gut sample, were in close proximity to the ISC1 cluster based on the UMAP coordinates, whereas Tk^+^EEs were distant from ISCs, suggesting transcriptional similarities between the AstC^+^EE subpopulation and ISCs in the early adult intestine (Figures 4A and 4B).

**Figure 4.**
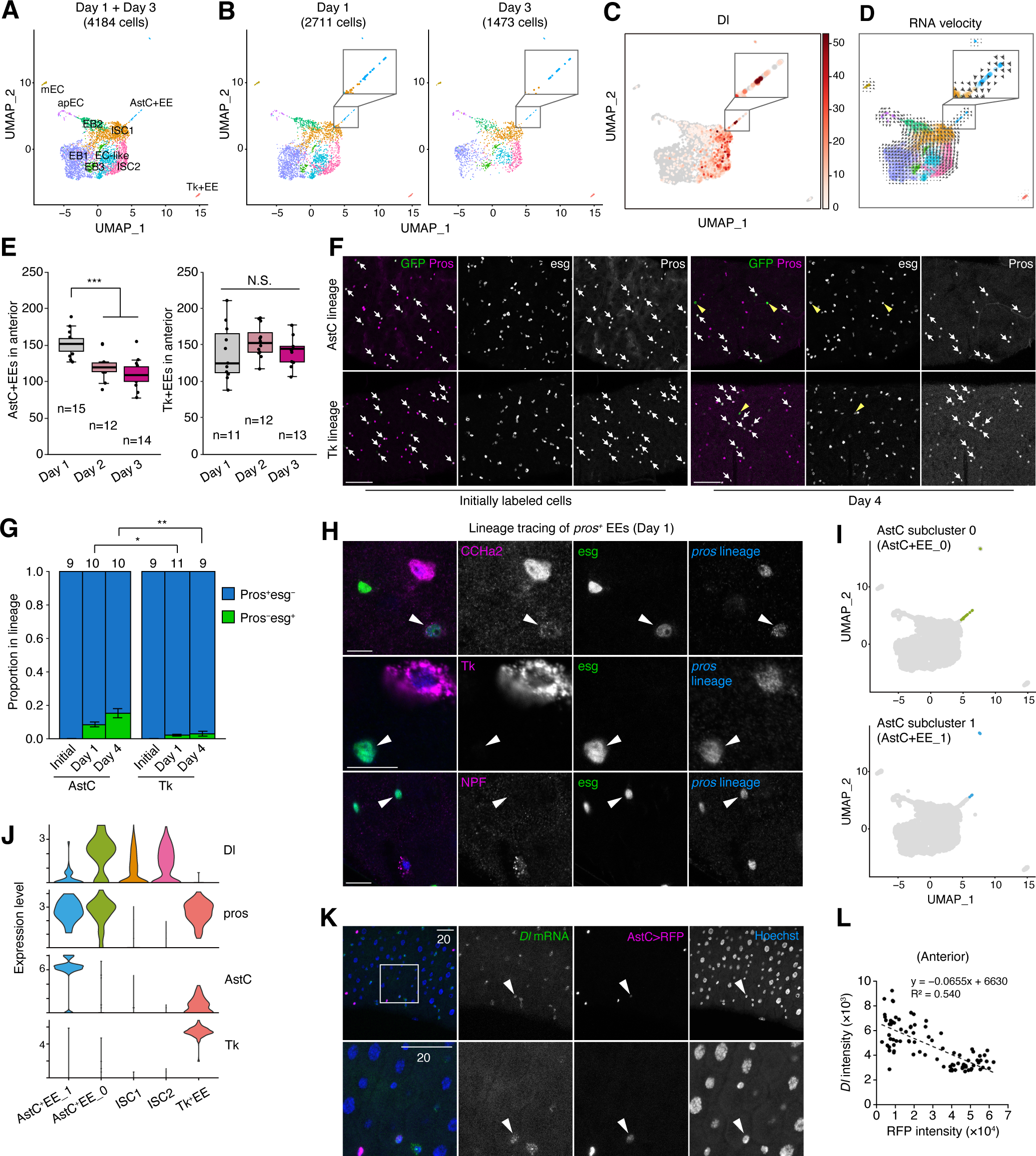
scRNA-seq identifies a subpopulation of EEs undergoing dedifferentiation. (A) UMAP projection of the 4,184 cells that passed quality control filtering. Data from Day 1 and Day 3 guts were merged and subjected to a graph-based clustering using the Louvain algorithm with Seurat v.4. (B) Side-by-side UMAP embedding showing the distribution of cells in Day 1 and Day 3 samples. (C) Projection of *Dl* mRNA levels onto the UMAP plot. (D) Projection of RNA velocities onto the UMAP plot. A subset of AstC^+^EEs exhibit direction toward the ISC1 cluster (inset). (E) The number of *AstC-Gal4>UAS-GFP^+^*cells and *Tk-Gal4>UAS-GFP^+^* cells in Day 1, 2, and 3 fed anterior midguts. (F) Representative images of AstC/Tk lineage tracing. Arrows: Pros^+^esg^−^ cells, arrowheads: Pros^−^esg^+^ cells. (G) Quantification of the Pros^+^esg^−^ and Pros^−^esg^+^ ratio in *AstC/Tk*-lineage cells. (H) Neuropeptide staining in the anterior region of *pros*-lineage tracing sample. In Day 1 fed guts, EE-derived esg*^+^* cells (arrowheads) contain remnants of CCHa2 peptide but not of Tk or NPF. (I) Projection of AstC^+^EE subclusters onto the UMAP plot. (J) Expression of the ISC marker (*Dl*) and the EE markers (*pros, AstC, Tk*) in the indicated cell population. (K) Representative image for SABER FISH of *Dl* mRNA in the *AstC-Gal4>UAS-RFP* midgut. (L) Quantification of (K). A correlation analysis of mean fluorescence intensity of *Dl* mRNA and *AstC>RFP* indicates that AstC^+^EEs exhibiting high *Dl* mRNA signal show low RFP signal, and vice versa. Pearson’ correlation coefficient (R) was calculated: R=−0.735, R^2^ = 0.540. N.S., not significant: P>0.05, * P≤0.05, **P≤0.01, ***P≤0.001. *n* indicates the number of midguts. One-way ANOVAs with post hoc Tukey test. Scale bars: 50 µm (F), 10 µm (H), 20 µm (K). See also Figure S4.

To identify EEs that undergo dedifferentiation, we next obtained RNA velocities and the directional information by performing trajectory inference analysis^54–56^. AstC^+^EEs exhibited direction toward ISC1 and ultimately ended in ISC2, while Tk^+^EEs had no specific direction toward other clusters (Figure 4D). Importantly, the number of AstC^+^EEs, but not Tk^+^EEs, decreased after eclosion *in vivo* (Figure 4E), and lineage tracing using *AstC-Gal4* or *Tk-Gal4* drivers confirmed that *AstC*-lineage more frequently converts to *esg^+^* cells than does *Tk*-lineage (Figures 4F and 4G). Consistent with these data, dedifferentiating EEs did not contain remnants of class II (Tk^+^) neuropeptides Tk or NPF, which was in stark contrast to the case of class I (AstC^+^) neuropeptides CCHa1/2 (Figures 3C, 4H, S4H)^46^.

Because RNA velocity analysis suggested that not all AstC^+^EEs have a direction toward ISCs, we further performed subclustering and identified two subpopulations identified as AstC^+^EE_0 and AstC^+^EE_1 (Figure S4I). AstC^+^EE_0 is formed by the majority of cells closer to ISC1 whereas AstC^+^EE_1 primarily constitutes the distant AstC^+^EE cells on the UMAP coordinates (Figure 4I). Integration with the previous scRNA-seq data from FACS-sorted EEs^46^ revealed that AstC+EE_0 represented Class I EEs in the anterior/posterior region that also showed similarity to ISCs, while AstC+EE_1 and Tk+EE represented EEs in the middle midgut (Figures S4J and S4K). Notably, AstC^+^EE_0 expressed both the ISC marker *Dl* and the EE marker *pros* while lowering transcription of the neuropeptide *AstC*, suggesting their intermediate state during dedifferentiation (Figure 4J). Consistently, we observed *AstC^+^Dl^+^* cells in the Day 1 anterior midgut, where the levels of *AstC* and *Dl* were inversely correlated (Figures 4K, 4L, and S4L). Furthermore, AstC^+^EE_0 highly expressed genes involved in stem cell maintenance, including the actin remodeling factor *chic* (the *Drosophila* homolog of Profilin)^57, 58^, which is consistent with the morphological transformation of dedifferentiating EEs (Figures 3A, 3B, S3I, and S4M). These data together identify a subpopulation of AstC^+^EEs that undergo dedifferentiation during midgut growth after eclosion.

### Genetic ablation of EE-derived stem cell population

ISC expansion in the early adult stage drives nutrient-dependent intestinal growth^3, 31, 32^, and our results indicated that EE dedifferentiation could be a critical driver of adaptive tissue growth in the anterior midgut by providing an additional ISC pool. To test this hypothesis, we developed a genetic ablation strategy that allows for the selective elimination of the EE*-*derived ISCs from the midgut. In brief, the Gal4/UAS system with temperature-sensitive Gal80 allows transient FLP expression in EEs under the *pros-Gal4*. FLP flips out the transcriptional repressor *tub-QS* in EEs, and then *esg-QF2*, which recapitulates its original *esg-Gal4* pattern (Figure S5A), induces expression of the pro-apoptotic gene *reaper* (*rpr*) in the EE*-*derived ISCs (Figure 5A). By transiently shifting pupae to restrictive temperature (29°C) before eclosion, this strategy enables selective ablation of ISCs that originate from EEs present at eclosion (Figure 5B). We confirmed the efficiency of our ablation paradigm by labeling EE-derived ISCs with GFP. While control GFP expression labeled diploid cells in both the anterior and posterior regions of the adult midgut, *rpr* expression together with GFP reduced GFP^+^ cells (Figures 5C, 5D, and S5B). Although *pros-Gal4* is active in neurons as well as in EEs, *pros*-derived *esg^+^* cells were not detected in the adult brain due to the lack of *esg-QF2* expression in neurons (Figures S5C and S5D). We can therefore conclude that genetic ablation occurs exclusively in EE-lineage cells in the midgut.

**Figure 5.**
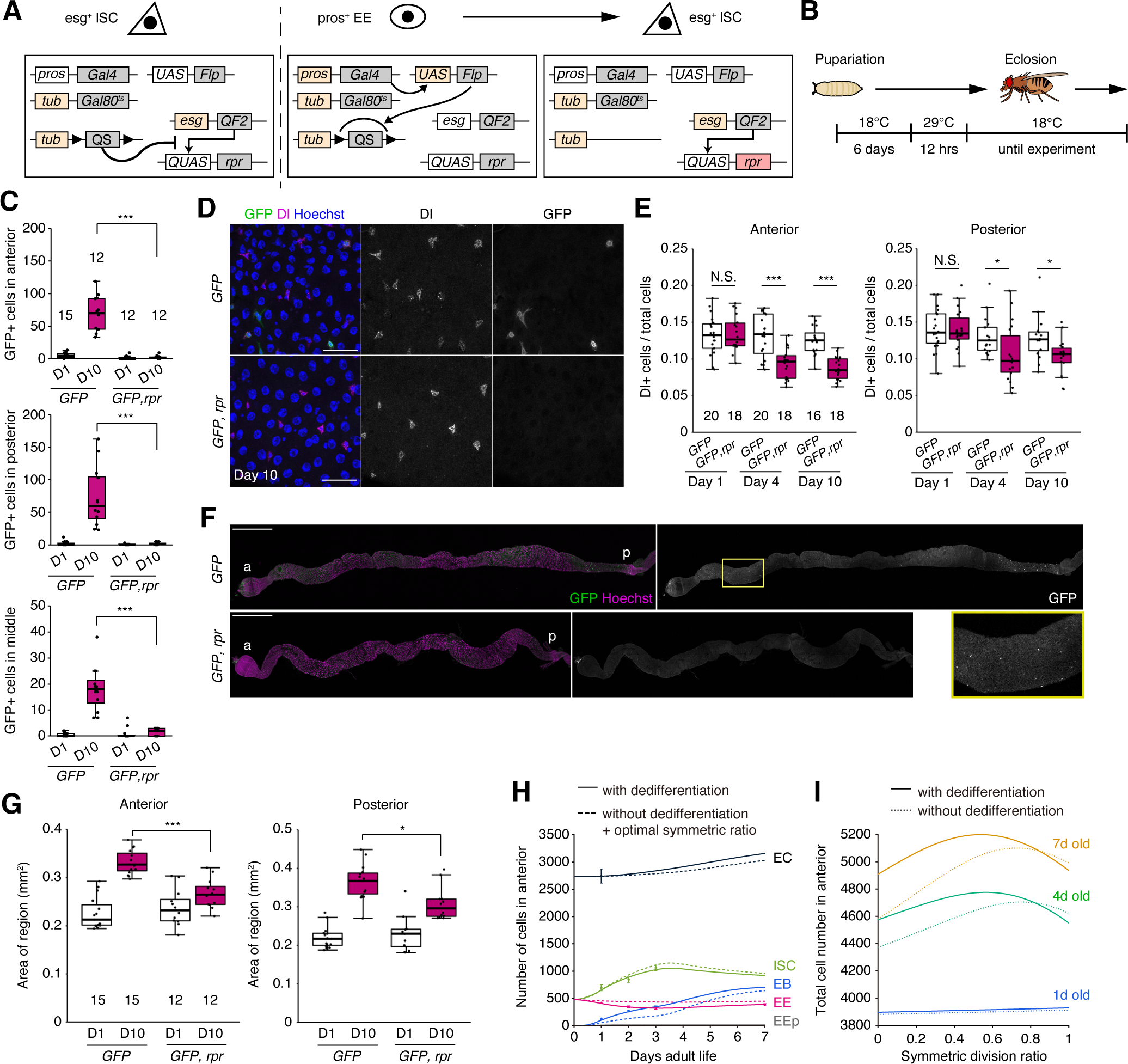
Dedifferentiation of EEs contributes to nutrient-dependent intestinal growth. (A) Schematic of the genetic system that allows ablation of EE-derived ISCs. (B) Ablation experiment scheme. (C) Ablation of *pros*-lineage *esg^+^* cells by *rpr* induction at Day 1 and Day 10. (D and E) Representative images of anti-Dl^+^ cells and EE-derived *esg-QF2>GFP^+^*cells in the control (*GFP*) and the ablated (*GFP, rpr*) anterior midguts at Day 10 (D). Dl^+^ cell abundance is quantified in (E). (F and G) Representative images of the control and the ablated whole midgut at Day 10. Size of the guts is quantified in (G). (H) Population dynamics in the anterior region over time. Two conditions, one wherein EEs undergo dedifferentiation and the other wherein ISCs divide more symmetrically due to the lack of EE dedifferentiation, are simulated. Dots and error bars (mean ± SE) indicate the cell number observed *in vivo*. (I) Computational simulation indicates the effect of symmetric ISC division on the total cell number in the anterior midgut with or without dedifferentiation. N.S., not significant: P>0.05, *P≤0.05, ***P≤0.001, two tailed *t* tests. *n* indicates the number of midguts in (C) and (G), and the number of images analyzed in (E). Scale bars: 10 µm (D), 500 µm (F). See also Figure S5 and S6.

### EE-to-ISC conversion contributes to nutrient-dependent midgut growth

Using the ablation system for EE-derived ISCs, we examined the impact of EE dedifferentiation on stem cell abundance in the adult midgut by measuring the proportion of Dl^+^ ISCs. After ablation of EE-derived stem cells, the Dl^+^ ISC ratio decreased significantly in Day 4 fed adults both in the anterior and posterior midgut with a stronger effect in the anterior region (Figures 5D and 5E), consistent with the higher frequency of dedifferentiation in the anterior midgut (Figures 2G and 3H). Surprisingly, the decreased Dl^+^ ratio persisted in Day 10 fed guts even though the priming of *rpr* induction was restricted exclusively to EEs existing at eclosion, suggesting that loss of EE-derived ISCs cannot be recovered via other mechanisms (Figure 5E). The decline in the Dl^+^ ISC ratio was not observed in either Day 4 starved adults or in Day 10 fed adults that did not experience the *rpr* induction priming (Figures S5E-S5H).

To determine if organ size increase requires EE dedifferentiation, we measured the size of adult midguts after ablation. The ablation of dedifferentiated ISCs significantly impaired organ growth in response to food intake after eclosion, particularly by attenuating the increase in thickness (Figure 5F, 5G, S5I, and S5J). Importantly, the reduction of organ growth was not caused by any abnormality in feeding behavior since *rpr* induction did not affect food intake (Figure S5K and S5L).

While the cell ablation experiments suggested that EE-to-ISC conversion provides an additional stem cell pool for efficient midgut growth, *rpr* induction ablated not only EE-derived ISCs in the anterior/posterior midgut but also Pros^+^esg^+^ EEs in the middle midgut (Figure 5C)^46, 48^. To eliminate any potential effect caused by the loss of middle EEs, we inhibited mitosis in the EE-derived ISCs by knocking down *cdk1, AurB,* or *polo* ^59^. After confirming that mitotic inhibition did not affect the Pros^+^esg^+^ EEs in the middle region (Figures S5M-S5O), we found that knockdown of these mitosis-related genes impaired growth of the anterior midgut, but not of the posterior midgut (Figure S5P). Therefore, the mitosis of EE-derived ISCs is the predominant contributor to the resizing of the anterior midgut.

Results from the cell ablation and mitotic inhibition experiments suggested that EE-to-ISC conversion provides an additional stem cell pool for efficient midgut growth. To further test this concept without blocking the functions of EE-derived ISCs, we established a population dynamics model that recapitulates our observations of cell population changes in the early adult midgut (Figures 5H [with dedifferentiation], S6, and Table S2). In this model, dedifferentiation occurs only during the first four days after eclosion, mirroring the life stage when the EE-to-ISC conversion occurs (Figures 2F and 2G). *In silico* simulation revealed that, if the anterior midgut does not rely on the dedifferentiation of EEs, ISCs must increase the proportion of symmetric self-renewing division to maximize the expansion of total cells (Figure 5I). The shift of division mode to symmetric division decreased the production of new ECs (Figure 5H). Intriguingly, the proportion of symmetric division in the anterior midgut *in vivo* (Figure 1E) was close to the optimal value (0.55) estimated by the mathematical model for increasing midgut cell number (Figure 5I). These results indicate that EE dedifferentiation functions as an irreplaceable source of new ISCs that relieves the need for symmetric ISC division and promotes the generation of new ECs. Consistently, the higher frequency of dedifferentiation in the anterior midgut (Figure 2G) accompanied a higher ratio of asymmetric ISC division at Day 2 and Day 3 compared to that in the posterior midgut *in vivo* (Figure 1E), further supporting the role of EE dedifferentiation in promoting EC generation.

### Dietary glucose and amino acids induce EE dedifferentiation

To gain insight into EE dedifferentiation mechanisms, we first investigated the nutrients required for the EE-to-ISC conversion by culturing lineage-tracing fly adults on holidic medium, a synthetic fly food^60^. Holidic medium lacking either sucrose or amino acids (AAs) significantly reduced the frequency of EE dedifferentiation, and food lacking both sucrose and AAs almost completely eliminated it to near the level of the water-only condition (Figure 6A). In contrast, dietary cholesterol was not necessary for EE dedifferentiation (Figure 6A). Intriguingly, ingestion of both sucrose and AAs induced cell fate conversion, albeit at a lower frequency than that induced by nutrient-complete medium (Figure 6B). These results suggest that dietary sugar and AAs are minimal nutrients required for dedifferentiation, while other nutrients also promote it.

**Figure 6.**
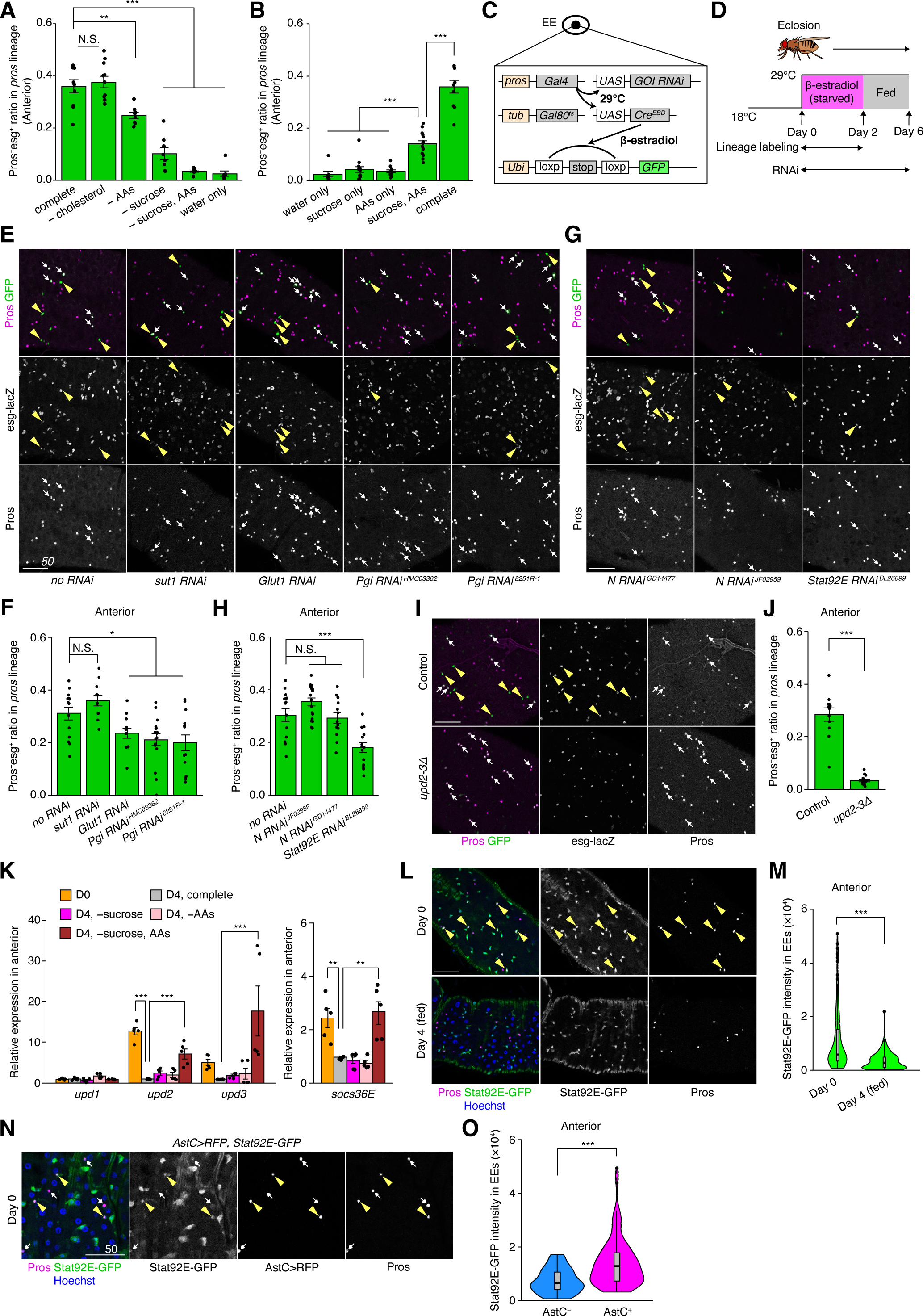
Dietary glucose and amino acids as well as the JAK-STAT pathway regulate EE dedifferentiation. (A, B) Quantification of the Pros^−^esg^+^ ratio in *pros-*lineage cells in the Day 4 anterior midgut. (C) Schematic of the T-trace system. (D) Scheme for the T-trace in the early adult stage. (E, G) Representative images for the T-trace of *pros* lineage in the anterior midgut. Arrows: Pros^+^esg^−^ cells, arrowheads: Pros^−^esg^+^ cells. (F, H) Quantification of Pros^−^esg^+^ ratio in *pros-*lineage cells in T-trace experiments. The Day 4 anterior midguts were analyzed. (I) Representative images for *pros*-lineage cells in the control and *upd2-3Δ* anterior midgut. Arrows: Pros^+^esg^−^ cells, arrowheads: Pros^−^esg^+^ cells. (J) Quantification of the Pros^−^esg^+^ ratio in *pros-*lineage cells in the Day 4 anterior midgut. (K) RT-qPCR for *upd1, upd2, upd3*, and *socs36E*. The anterior midguts were collected from Day 0 (D0) and Day 4 (D4) adults. (L) Representative images of 10×Stat92E-GFP signal in the anterior midgut. Arrowheads: GFP^high^Pros^+^ cells. (M) Quantification of 10×Stat92E-GFP signal intensity in Pros^+^EEs. (N) Representative images of 10×Stat92E-GFP signal in AstC-Gal4>RFP anterior midgut. Arrowheads: GFP^high^Pros^+^ cells, arrows: GFP^low^Pros^+^ cells. (O) Quantification of 10×Stat92E-GFP signal intensity in AstC^+^Pros^+^EEs and AstC^−^Pros^+^EEs in the Day 0 anterior midgut. N.S., not significant: P>0.05, * P≤0.05, **P≤0.01, ***P≤0.001. One-way ANOVAs with post hoc Tukey test. *n* indicates the number of guts (A, B, F, H, J), RNA samples (K), and cells (M, O). Scale bars: 50 µm. See also Figure S7.

The feeding assay used the fluorescently labeled deoxyglucose (2-NBDG) revealed that anterior EEs incorporated more glucose than posterior EEs did (Figure S7A-S7B), raising the possibility that glucose directly acts on EEs to regulate their plasticity. To test this hypothesis, we introduced another lineage tracing system, T-trace^61, 62^, in which lineage labeling requires not only temperature shift but also estrogen feeding (Figure 6C). This two-step regulation enables us to knock down genes of interest in EEs while performing lineage tracing (Figures 6C and 6D). We first confirmed that T-trace exhibited no leaky labeling during our tracing duration and reproduced the regional difference in the frequency of EE-to-ISC conversion (Figures S7C-S7E). Then we tested the requirement of two glucose transporters, Glut1 and Sut1, which have been reported to function in EEs^63, 64^, as well as Pgi, a downstream glycolytic enzyme. Knockdown of *Glut1* and *Pgi*, but not *sut1*, suppressed cell fate conversion (Figures 6E and 6F). Moreover, the *Pgi:GFP* reporter^65^ revealed that anterior EEs expressed more Pgi protein than posterior EEs in Day 1 midguts (Figures S7F and S7G). These results suggest that EEs directly sense glucose and metabolize it to revert into stem cells.

### The JAK-STAT pathway underlies EE-to-ISC conversion

Given that several signaling pathways (e.g. Wnt, Notch, and EGFR) have been reported to regulate cellular reprogramming during intestinal regeneration^19, 23, 24, 66^, we next performed candidate screening to identify the signaling pathway underlying the nutrient-dependent dedifferentiation of EEs. In this screening, we repressed signaling factors in EEs using *pros-Gal4* and counted the number of Pros^+^EEs at Day 3, when EEs decreased in the control midgut (Figures S2A and S2B). Knockdown of *Notch, Stat92E,* and *domeless* (a receptor in the JAK-STAT pathway) resulted in a significant increase of EEs (Figures S7H-S7L). From T-trace experiments, we identified *Stat92E*, but not *Notch*, as a regulator of EE-to-ISC conversion (Figure 6G and 6H). Furthermore, flies lacking both *upd2* and *upd3* (*upd2-3Δ*), which encode ligands for the Domeless receptor, failed to induce the dedifferentiation (Figure 6I and 6J). These results indicate that the JAK-STAT pathway is crucial for the cell fate reversion of EEs.

Previous work showed that starvation induces *upd3* expression in the adult midgut^67^, raising a possibility that the JAK-STAT pathway is activated during food scarcity. Indeed, the expression of *upd2, upd3*, and *socs36E* (a downstream target of Stat92E), but not *upd1*, was high in pre-feeding Day 0 (“D0”) anterior midguts, but their expression decreased after food intake (“D4, complete”) (Figure 6K). When dietary sucrose and AAs were depleted from fly food, the levels of *upd2, upd3*, and *socs36E* remained high in Day 4 anterior midguts (Figure 6K), suggesting that the JAK-STAT pathway continues to be activated until adult flies ingest enough nutrients to induce dedifferentiation (Figure 6A). We further found that transcriptional activity of Stat92E was high in Day 0 EEs compared to EEs in the Day 4 fed condition (Figures 6L and 6M). Importantly, AstC^+^EEs exhibited higher Stat92E activity among Pros^+^ population (Figure 6N and 6O), and in scRNA-seq data, AstC+EE_0 expressed *domeless, Stat92E, and Socs36E* to a higher degree than Tk+EE (Figure S4N), which is in line with the higher plasticity in this EE subtype (Figure 4D and 4G). The *upd3-Gal4>GFP* reporter also revealed that *upd3* was upregulated in the Day 0 midguts (Figures S7M and S7N). Consistent with the previous report^67^, it was not EEs but mainly ECs that produced *upd3* in the anterior midgut (Figure S7O). Collectively, Stat92E is activated in anterior EEs under nutrient-restricted conditions, which is necessary to induce dedifferentiation in response to subsequent food intake.

### Dedifferentiation of EEs occurs generally in response to nutrient fluctuation

Given that the midgut of the newly eclosed adult is food-naïve due to the lack of food intake during the pupal stages, fluctuation in nutrient conditions may trigger fate conversion of EEs throughout life. To test this hypothesis, we examined the behavior of EEs after a feed-starve-refeed cycle and found that the total cell number increased in response to refeeding^3^ (Figures 7A, S7P, and S7Q). The number of EEs, measured using *pros-Gal4* (Figure 7B) or anti-Pros (Figure 7C), significantly decreased upon refeeding in the anterior midgut, suggesting that anterior EEs maintain the potential to dedifferentiate even after midgut maturation. Concordantly, lineage tracing revealed that EEs, especially AstC^+^EEs, convert into *esg^+^* cells after refeeding in the anterior region (Figures 7D and 7E). The behaviors of the EE-derived *esg^+^* cells were similar to those in the early adult midgut: after 7 days of refeeding, they clonally expanded and exhibited multipotency as well as differentiation bias toward ECs, although the ratio of the EC-only clones was lower compared to that in the early adult midgut (Figures 7F-7J). Moreover, *Stat92E* is required in EEs to induce the EE-to-ISC conversion, and the transcriptional activity of Stat92E was high in AstC^+^EEs compared to other EEs before refeeding (Figures 7K-7N). Taken together, these results indicate that dedifferentiation of EEs can occur generally during recovery from starvation.

**Figure 7.**
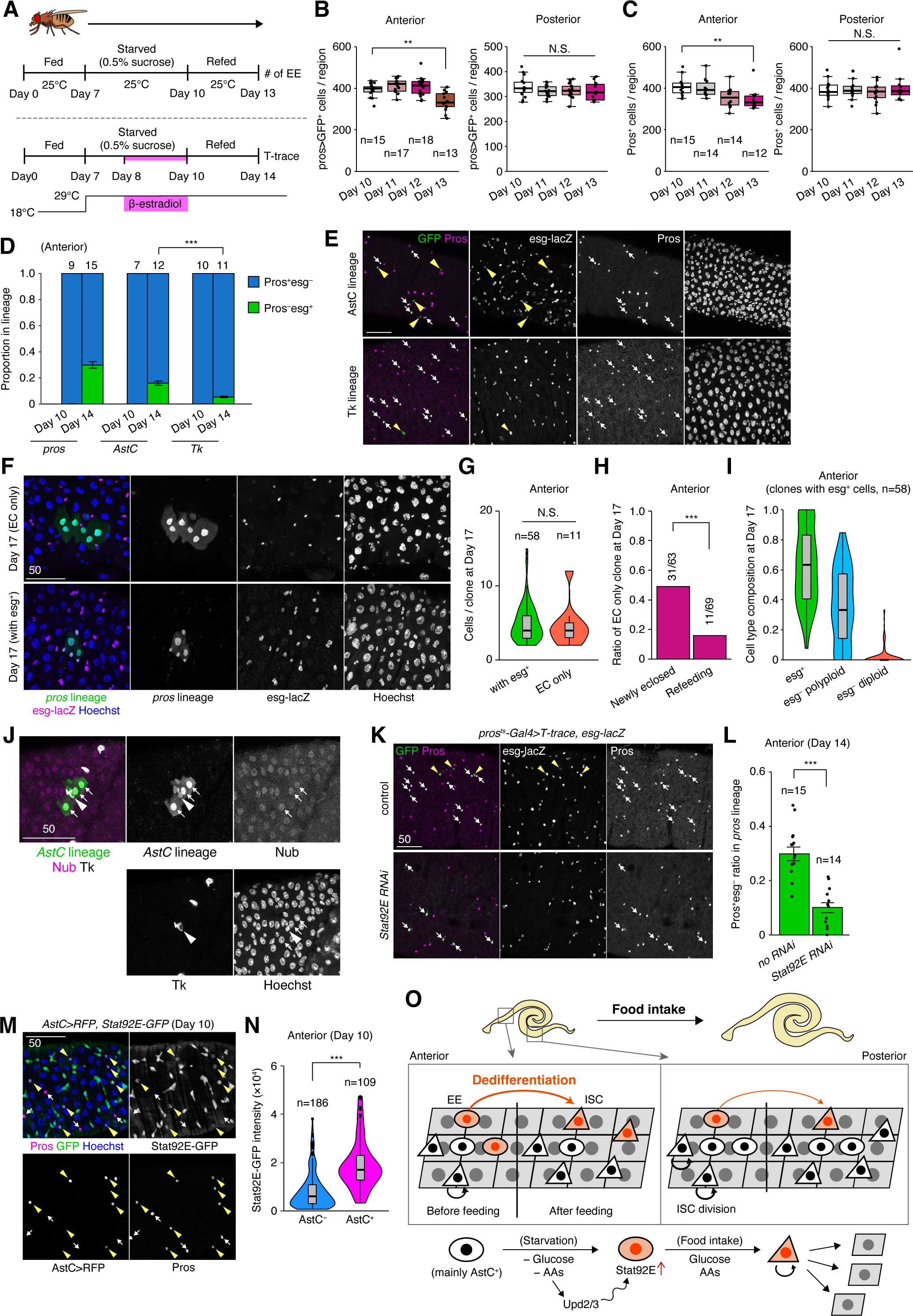
Dedifferentiation of EEs occurs generally in response to nutrient fluctuation. (A) Experimental schematic of the feed-starve-refeed cycle. Newly eclosed female adults were fed for 7 days, starved for 3 days (0.5% sucrose), and then refed for 1, 2, 3, or 4 days. Lineage labeling was performed during the last two days of starvation (from Day 8 to Day 10). (B, C) Refeeding decreased the number of *pros>GFP^+^* cells (B) and anti-Pros^+^ cells (C) in the anterior midgut. No decrease was observed in the posterior midgut. (D) Quantification of the Pros^+^esg^−^:Pros^−^esg^+^ ratio in *pros/AstC/Tk*-lineage cells in the anterior midgut. (E) Representative images of *AstC/Tk* lineage tracing in the Day 14 anterior midgut. Arrows: Pros^+^esg^−^ cells, arrowheads: Pros^−^esg^+^ cells. (F) Representative images of *pros*-lineage clones 7-days after refeeding (Day 17). The clone containing only polyploid ECs (EC only) and the one retaining esg^+^ cells (with esg^+^) are shown. (G) The number of cells per *pros*-lineage clone at Day 17 in the anterior midgut. (H) The ratio of EC-only clones in *pros*-lineage clones. (I) The cell type composition in *pros*-lineage clones that retained esg^+^ cells at Day 17. (J) Nub^+^ECs and a Tk^+^EE in one clone that derived from an AstC^+^EE. Arrows: Nub^+^ECs, arrowhead: Tk^+^EE. (K) Representative images of *pros*-lineage cells in the control and *Stat92E RNAi* midgut. (L) Quantification of the Pros^−^esg^+^ ratio in *pros-*lineage cells. (M) Representative images of 10×Stat92E-GFP signal in the *AstC-Gal4>RFP* anterior midgut. Arrowheads: GFP^high^Pros^+^ cells, arrows: GFP^low^Pros^+^ cells. (N) Quantification of 10×Stat92E-GFP signal intensity in Pros^+^ cells. (O) Model schematic. The anterior midgut highly relies on EE dedifferentiation for nutrient-dependent intestinal growth, whereas symmetric ISC division is the dominant mechanism in the posterior midgut. The EE-to-ISC conversion is regulated by dietary glucose and AAs as well as the JAK-STAT pathway. N.S., not significant: P>0.05, **P≤0.01, ***P≤0.001. Two-tailed *t* tests (D, G, L, N), chi- square test (H). *n* indicates the number of midguts (B, C, D, L), the number of clones observed (G-I), and the number of cells (N). See also Figure S7.

## DISCUSSION

Here, we demonstrate that dedifferentiation of EEs occurs during adaptive midgut resizing when the number of ISCs additively increases in early *Drosophila* adults (Figure 7O). Although cell fate plasticity *in vivo* has been well documented under the conditions of stem cell loss, our results provide evidence that physiologically-induced dedifferentiation contributes more broadly to stem cell expansion beyond the cases of regeneration and disease. Indeed, enteroendocrine lineage in the mice intestine exhibits rare stem cell activity even without severe injury^26^. Given that diverse species including mammals dynamically resize digestive organs depending on nutrient availability^2, 4–6^, it is tempting to speculate that dedifferentiation is an evolutionarily conserved mechanism underlying adaptive tissue growth.

Both in mammals and flies, EEs are diversified according to neuropeptide expression, and specific subtypes sense different types of luminal environment to induce local and/or systemic responses^68, 69^. In *Drosophila*, class II EEs secrete Tk, which activates ISC proliferation via insulin signaling upon food intake^30^. The higher cell fate plasticity of a subset of class I EEs (Figures 4F and 4G) whose endocrine functions are more important during starvation than under fed conditions^70, 71^ likely indicates that paracrine signaling from class II EEs and dedifferentiation from class I EEs cooperatively promote ISC expansion in response to food intake. Although dedifferentiation causes a partial loss of class I EEs, the supply can be restored after intestinal growth (Figures 2A and S2A), suggesting that the enlarged intestine replenishes the starvation-responsive population to prepare for potential future food scarcity.

While nutritional inputs shift the division mode of ISCs toward symmetric renewal^3, 31, 32, 72^, the mechanisms employed to sustain the generation of differentiated cells during midgut growth are unclear. Our mathematical modeling shows that the existence of nutrient-dependent EE dedifferentiation secures EC lineage-generating asymmetric ISC division by supplying EE*-*derived ISCs during the rapid midgut growth phase (Figures 5H and 5I), highlighting the potential significance of physiological dedifferentiation for organ growth. Moreover, the EE-derived ISC itself also preferentially generates ECs, especially in the anterior midgut (Figures 3G, 3I, 7F and 7H). Given the critical roles of ECs in digestion and absorption, the generation of new ECs in the growing intestine may help to optimize the intestine’s capacity to maximize nutrient availability. This digestive function is particularly important in the anterior region, a major site of macromolecule degradation essential for subsequent nutrient absorption^9, 11^. Consistent with this notion, the *Drosophila* anterior midgut exhibits a higher dedifferentiation rate with a relatively high ratio of asymmetric ISC division (Figures 1E and 2G).

While cell fate reversion during intestinal regeneration relies on Wnt, Notch, and EGFR pathways^19, 23, 24, 66^, our candidate screening newly identified Stat92E as a signaling factor required for the nutrient-dependent dedifferentiation of EEs (Figures S7H-S7L, 6G, and 6H). In line with our finding, the ligands of the JAK-STAT pathway, but not those of Wnt and EGFR pathways, are specifically up-regulated in the adult midgut during starvation^67^. Interestingly, activated STAT3 binds to progenitor-related genes to induce dedifferentiation of mouse hepatocytes during liver regeneration^73^. However, in the case of nutrient-dependent intestinal growth, refeeding of glucose and AAs is also required in addition to the Stat92E activity to trigger the dedifferentiation process (Figures 6A and 6B). Future studies should investigate how dietary glucose and AAs cooperate with Stat92E to induce EE-to-ISC conversion in response to refeeding.

On the basis of our findings, we propose that the coordination of cell fate plasticity and stem cell division ensures functional organ growth in which both stem cells and differentiated cells concomitantly increase their number in response to nutrition changes. In this scenario, EEs may enable an “on-demand” supply of additional ISCs by sensing luminal nutrients^68, 69^, which fluctuate with food availability in the wild as well as under pathophysiological conditions^74, 75^. The number of EEs remains constant during starvation (Figure 2B and S2B), supporting the idea that EEs function as a backup population that undergoes dedifferentiation only when responding to tissue demand for stem cells. Collectively, our study illuminates the physiological regulation of cell fate plasticity and its role in adaptive organ resizing.

### Limitations of study

In this study, we investigated the cell fate plasticity that underlies the nutrient-dependent intestinal growth. Although intestinal size can dynamically change under other physiological contexts such as mating^76–79^ and regeneration^80, 81^, it remains to be investigated whether these external stimuli also induce cell fate reversion of EEs. Notably, it was reported that pathogenic infection by *Pseudomonas entomophila* did not alter the identity of either Class I (AstA^+^) EEs or Class II (Tk^+^) EEs^82^, while EBs revert into ISCs in response to the bacterial infection^66^. It is thus possible that the cell type undergoing dedifferentiation may vary with physiological context. DSS-induced enteritis induces reversion of Paneth cells in the mouse intestine^23^, raising the possibility that inflammatory cues, including Upd3 (orthologous to mammalian IL-6), identified in this study, regulates cellular reprogramming during inflammation. Consistently, macrophage-derived IL-6 induces dedifferentiation of hepatocytes during liver regeneration^73^. It will be worthwhile to investigate whether nutritional and Stat-dependent mechanisms control cell fate reversion beyond starvation-refeeding contexts.

## ACKNOWLEDGEMENTS

We thank H. Jasper, N. Perrimon, T. Akiyama, H. Bruce, H. Steller, S. Bray, R. Niwa, T. Ida, E.Y. Kim, N. Buchon, S. Hou, I. Miguel-Aliaga, BDSC, the Kyoto Stock Center, NIG, and DSHB for fly stocks and reagents; S. Kondo and T. Akiyama for critical suggestions for and comments on the manuscript.

This work was supported by JSPS/MEXT KAKENHI (grant numbers JP22J01430 to H.N., JP17H06327, JP19K03645 to S.T., JP17H06331 to R.N., JP21H05105 to D.U., JP26114003, JP21H05255, JP24687027, JP16H04800 to E.K., JP21H04774, JP21K19206 to M.M., and JP17H06332, JP19K22550, JP22H02762 to Y.N.), JST CREST (JPMJCR1852 to E.K.), JST FOREST Program (J210000474 to D.U.), AMED-Aging (JP21gm5010001 to M.M.), AMED-PRIME (JP22gm6310012 to R.N., JP21gm6110025 to Y.N.), Takeda Science Foundation (D.U.), and Sadako O. Hirai Ban Award for Young Researchers (H.N.).

## AUTHOR CONTRIBUTIONS

Conceptualization: H.N., Y.N. Investigation: H.N., L.A.E.N., S.T.

Methodology: H.N., L.A.E.N., S.T., D.U.

Validation: H.N., L.A.E.N., S.T., D.U., E.K., R.N., M.M., Y.N.

Data curation: H.N., L.A.E.N., S.T., Y.N.

Writing – original draft: H.N., L.A.E.N., S.T., D.U.

Writing – review & editing: H.N., L.A.E.N., S.T., D.U., E.K., R.N., M.M., Y.N.

Supervision: E.K., R.N., M.M., Y.N.

Funding acquisition: H.N., S.T., R.N., D.U., E.K., M.M., Y.N.

## STAR Methods

### RESOURCE AVAILABILITY

#### Lead contact

Further information and requests for resources and reagents should be directed to and will be fulfilled by the lead contact, Yu-ichiro Nakajima.

#### Materials availability

All *Drosophila* stocks generated in this study are available from the Lead Contact without restriction.

#### Data and code availability

- Raw scRNA-seq datasets are available from NCBI GEO (accession number GSE207662). Microscopy data reported in this paper will be shared by the lead contact upon request.
- The docker image used in the single-cell analysis is available at DockerHub (https://hub.docker.com/r/rnakato/shortcake). The scRNA-seq analysis scripts are available on GitHub (https://github.com/eijynagai/Drosophila_dedifferentiation_Nagai).
- Any additional information required to reanalyze the data reported in this paper is available from the lead contact upon request.

### EXPERIMENTAL MODEL AND SUBJECT DETAILS

#### Drosophila stocks

All stocks were maintained on a standard diet containing 4% cornmeal, 6% baker’s yeast (Saf Yeast), 6% glucose (Wako, 049-31177), and 0.8% agar (Kishida chemical, 260- 01705) with 0.3% propionic acid (Tokyo Chemical Industry, P0500) and 0.05% nipagin (Wako, 132-02635). Canton S was utilized as the wild type strain. Transgenic fly lines were obtained from Bloomington *Drosophila* Stock Center, Kyoto Stock Center, NIG-FLY, Vienna *Drosophila* Resource Center. Following lines are gifts from fly community: *w; esg-Gal4, UAS-eYFP; tub-Gal80ts, Su(H)GBE-Gal80* (Deng et al., 2015)^83^, *w; Dl- Gal4* (Zeng et al., 2010)^84^, *w; upd3-Gal4* (Agaisse et al., 2003)^85^, *UAS-myc::DIAP1* (Hay et al., 1995)^86^, *yw;; QUAS-rpr* (Pérez-Garijo et al., 2013)^87^,. *w;; UAS-FLP, Act-FRT-stop-FRT-lacZ* (Akiyama and Gibson, 2015)^88^, *w; Ubi-loxP-stop-loxP-GFP* (Zeng and Hou, 2015)^61^, *w;; tub-Gal80ts, UAS-Cre[EBD304]* (Zeng and Hou, 2015)^61^, *esg- GFP[P01986]* (Le Bras and Van Doren, 2006)^89^, *Su(H)GBE-lacZ* (Furriols and Bray, 2001)^90^, *w; Pgi:GFP* (Hudry et al., 2019)^65^. Following lines are generated in this study: *w; esg-QF2*, *w;; QUAS-Cdk1 RNAi*, *w;; QUAS-AurB RNAi*, *w;; QUAS-polo RNAi*, *w; Myo31DF-Venus.* See **Table S3** for the genotypes present in each figure.

### METHOD DETAILS

#### *Drosophila* genetics

Virgin female adults were used in all experiments. When Day 0 adults were raised under starvation, raised on holidic medium, and treated with 2-NBDGs, female adults were collected within 2 hours after eclosion.

Experimental crosses that did not involve Gal80^ts^-mediated temporal control were performed at 25°C. When using Gal80ts, experimental crosses were maintained at 18°C, and female white pupae were transferred to new vials. The collected pupae were raised at 18°C and then shifted to 29°C per the following time course: 18°C for 7 days and then 29°C for 1, 2, or 3 days (**Figure 1B, 1C, 1F, S1B, S2G, and S2H**); 18°C for 6 days, 29°C for 12 hrs, and then 18°C until experiments (**Figure 2F, 2G, 3A-3E, 4F-4H, 5C-5G, 6A, 6B, 6I, 6J, S3E-S3J, S3O, S4H, S5B, S5D, S5E, S5G, S5I-S5K, and S5P**); 18°C for 6 days, 29°C for 2 hrs, and then 18°C until experiments (sparse labeling, **Figure 3G-3K and S3K-S3N**).

In T-trace experiments in the early adult stage (**Figure 6E-6H and S7D-S7E**), Day 0 adults were transferred to 29°C and fed with 300 µg/ml β-estradiol (Sigma, E4389) dissolved in 0.5% (w/v) sucrose (Wako, 196-00015) for 2 days. Then flies were transferred to 18°C and fed with normal cornmeal food that did not contain β-estradiol for 4 days. In T-trace experiments in the feed-starve-refeed contexts (**Figure 7A, 7D, 7E, 7K and 7L**), female adults were fed for 7 days at 18°C, then starved by treating 0.5% sucrose for 3 days at 29°C. During the last two days of starvation, they were treated with 300 µg/ml β-estradiol. After starvation, flies were refed for 4 or 7 days at 18°C. For sparse labeling (**Figure 7F-7J**), 150 µg/ml β-estradiol (Sigma, E4389) was used.

For twin-spot MARCM analysis (**Figure 1D-1E and S1H**), female adults were collected within 2 hours after eclosion and maintained at 25°C for 1, 2, or 3 days. Then twin spot clones were induced by heat shock at 37°C for 1 hour. Symmetric or asymmetric outcome of the induced clones was determined 3-4 days after heat shock.

In the experiments for **Figure S5N**, 3-4 days old female adults were fed with 83 mg/ml quinic acid (Sigma, 138622, dissolved in 5% sucrose) at 18°C for 7 days to induce QF2-mediated knockdown of *cdk1, AurB*, and *polo*. We added 200 µl of the quinic acid solution on the top of the cornmeal food and put filter paper (Whatmann 3MM) on it to soak the solution.

#### Starvation experiments

For newly eclosed adults (**Figure 2B, 2G, S1F, S2B, S5E, S5G**), virgin females were collected within 2 hours after eclosion and transferred to vials with filter paper (Whatmann 3MM) soaked with 400 µl of water. For mature adults (**Figure 7**), 0.5% (w/v) sucrose was used instead of water. Flies were transferred to new vials every day during starvation.

#### Holidic medium

We followed the published recipe^60^ with modification based on exome matching^91^. The final concentration for each ingredient is: 15 g/L agar, 3g/L KH_2_PO_4_, 1g/L NaHCO_3_, 83.9 mg/L CaCl_2_·6H_2_O, 1.25 mg/L CuSO_4_·5H_2_O, 12.5 mg/L FeSO_4_·7H_2_O, 256 mg/L MgSO_4_·7H_2_O, 0.5 mg/L MnCl_2_·4H_2_O, 12.5 mg/L ZnSO_4_·7H_2_O, 0.3 g/L cholesterol, 17.2 g/L sucrose, 1.97 g/L L-arginine monohydrochloride, 0.65 g/L L-histidine, 1.71 g/L L-lysine monohydrochloride, 0.6 g/L L-methionine, 1.01 g/L L-phenylalanine, 1.11 g/L L-threonine, 0.32 g/L L-tryptophan, 1.2 g/L L-valine, 1.1 g/L L-alanine, 1.03 g/L L-asparagine, 1.52 g/L L-aspartic acid sodium salt monohydrate, 0.44 g/L L-Cysteine, 1.12 g/L L-Glutamine, 0.77 g/L Glycine, 0.98 g/L L-proline, 1.38 g/L L-serine, 1.75 g/L L-glutamic acid monosodium salt hydrate, 1.12 g/L L-isoleucine, 2.03 g/L L-leucine, 0.93 g/L L-tyrosine, 1.4 mg/L thiamine hydrochloride, 0.704 mg/L (−)-riboflavin, 8.45 mg/L nicotinic acid, 10.9 mg/L D-pantothenic acid hemicalcium, 1.76 mg/L pyridoxine hydrochloride, 0.14 mg/L biotin, 0.5 mg/L folic acid, 50 mg/L choline chloride, 5.04 mg/L myo-inositol, 65 mg/L inosine, 60 mg/L uridine, 6 ml/L propionic acid, and 10 ml/L nipagin.

#### Generation of *esg-QF2* line

The *esg-QF2* line was generated using the homology assisted CRISPR knock-in (HACK) method^92^, which converts the *X-Gal4* transgene into *X-QF2* through CRISPR-mediated introduction of double strand break and subsequent homology-directed repair. In brief, *esg-Gal4* (Kyoto Stock Center 104863) was crossed with *nos-Cas9*, and F1 embryos were injected with a *pBPGUw-HACK-G4>QF2* donor plasmid (Addgene #80277). Successful knock-in events were screened by identifying *w^+^*marker and eye marker *3*×*P3-RFP*. Injection and selection were performed by WellGenetics (Taiwan, R.O.C.).

#### Generation of *QUAS-cdk1/AurB/polo RNAi* line

To construct the *QUAS-cdk1, AurB, polo shRNA* plasmids, *pQUAS-WALIUM20* vector (*Drosophila* Genomics Resource Center, #1474) was digested with *Eco*RI and *Nhe*I, and then ligated with a DNA fragment for each gene (See **Table S4** for the sequences), based on pre-existing RNAi sequences (*cdk1*: HMS01531, *AurB*: HMJ22415, *polo*: HMS00530). The ligated plasmids were injected into *y[1] M{vas-int.Dm}ZH 2A w[*];P{y[+t7.7]=CaryP}attP2* embryos. Injection and selection were performed by WellGenetics (Taiwan, R.O.C.).

#### Generation of *Myo31DF-Venus* line

For the *Myo31DF* knock-in construct plasmid, the pBlueScriptII SK+ vector was digested with *Eco*RI, and then ligated with a cassette containing the fluorescent protein Venus whose sequence was excised from the pPVxRF3 plasmid with *Esp*3I and homologous recombination (HR) arms by the In-Fusion HD kit (Clontech). HR arms were amplified by PCR from genomic DNA extracted from a single CAS-0003 (NIG-FLY) adult fly. The knock-in construct was designed to insert the knock-in cassette containing the full length Venus sequence into the site in front of the termination codon of the gene. PCRs were performed using the primers 5’- GCTTGATATCGAATTCACAAGCAGGCTAATCGCGCCTTCATCG-3’ and 5’- AGTTGGGGGCGTAGGAACGCAGTACGCCGCCGGCACCTCG-3’ for the left HR arm and 5’-TAGTATAGGAACTTCGCGGAATCAACTCCGCCCAACTGTATTG-3’ and 5’-CGGGCTGCAGGAATTCTTTGGGGGAATTCATGACGAAATGACCG-3’ for the right HR arm. To construct the gRNA plasmid for CRISPR/Cas9, the pBFv-U6.2 vector was digested with BbsI and ligated with the double stranded oligo DNA sequences 5’-CTTCGCCTAAACGCAGTACGCCGC-3’ and 5’-AAACGCGGCGTACTGCGTTTAGGC-3’. To generate knock-in strains using CRISPR/Cas9, the gRNA plasmid and the knock-in plasmid were injected into the nos- Cas9 flies (CAS-0003 from NIG-FLY) as early embryos. The injection was performed by BestGene Inc. Isogenized DsRed-positive transformants were confirmed by genomic PCR and sequencing.

#### Immunofluorescence

Samples were dissected in 1X PBS and fixed in 4% PFA for 30-45 minutes at room temperature (RT). The following primary antibodies were used with indicated dilution into 1X PBS containing 0.5% BSA and 0.1% Triton X-100: rabbit anti-PH3 (Millipore 06-570, 1:1000), mouse anti-Prospero (DSHB MR1A, 1:100), rabbit anti-GFP (MBL 598, 1:500), rat anti-GFP (Nacalai tesque 04404-26, 1:400), rabbit anti-dsRed (Clontech 632496, 1:1000), chicken anti-β-galactosidase (Abcam ab9361, 1:500), mouse anti-Armadillo (DSHB N27A1, 1:100), rabbit anti-CCHa1 (T. Ida, 1:1000)^93^, rabbit anti- CCHa2 (T. Ida, 1:1000)^93^, guinea pig anti-NPF (R. Niwa, 1:2000)^64^, guinea pig anti-DTk (E.Y. Kim, 1:200)^94^, mouse anti-Nubbin (DSHB 2D4, 1:100), mouse anti-rCD2 (BIO-RAD MCA154GA, 1:2000), and mouse anti-Delta (DSHB C594.9B, 1:100).

After overnight incubation with primary antibodies at 4°C, samples were incubated with fluorescent secondary antibodies (Jackson ImmunoResearch and Invitrogen, 1:500) for 1 hour at RT. Hoechst 33342 (Invitrogen, final concentration: 10 µg/ml) was used to visualize DNA. Samples were mounted in Slowfade Diamond (ThermoFisher, S36963) and imaged with confocal microscopy (Zeiss LSM880 or Leica SP5). Whole midgut/brain images were obtained using the tile scan tool together with the z-stack tool (**Figure 1A, 5F, S3E, S5B-S5D, S7C, S7M, S7Q**). Other magnified images were taken from the R2 region of the anterior midgut unless noted otherwise in the figures.

#### TUNEL staining

Dissected midguts were fixed in 4% PFA for 1 hour at RT. The samples were then incubated with TUNEL reagents (Roche, 12156792910) in the dark at 37°C for 2 hours with 300 rpm shaking. The TUNEL signal was detected after Hoechst staining using the 543 nm He-Ne laser of the Leica SP5. For a positive control that increases TUNEL^+^ cells, we prepared flies that were fed with 5 mM paraquat (Sigma, 856177) overnight.

#### Sytox staining

Dissected midguts were incubated with 1 µM Sytox orange (Invitrogen, S11368) together with 10 µg/ml Hoechst33342 for 10 minutes at RT without fixation. The samples were then immediately observed with the Leica SP5. Paraquat was used for the positive control, as described in TUNEL staining.

#### Sample preparation for scRNA-seq

Digestive tracts were dissected in sterilized cold 1× PBS and stored on ice. We removed the hindgut, Malpighian tubules, and proventriculus to collect midguts (∼160 midguts for the Day 1 sample and ∼130 midguts for the Day 3 sample) after all samples were dissected. Midguts were then dissociated in 500 µl of 0.5% Trypsin-EDTA (Wako, 208-17251) at RT for 30 minutes with gentle pipetting every 10 minutes. The digestion was stopped by adding an equal amount of 1% BSA (Wako, 012-23381). Dissociated cells were passed through a 37 µm cell strainer, pelleted at 400 × g for 10 minutes at 4°C, and resuspended in 1% BSA. Cell suspension was loaded on the top of 1.12 g/ml gradient Optiprep reagent (Axis-Shield, 1114542). After centrifugation at 800 × g for 20 minutes, viable cells were isolated from the interphase, pelleted at 500 × g for 5 minutes, and resuspended in 100 µl of 0.1% BSA. Cell concentration and viability was assessed using auto cell-counter TC- 20 (BIO-RAD, 1450109J1) and 0.4% Trypan-blue (Wako, 207-17081). The samples (Day 1: 922 cells/µl, 81% viability; Day 3: 780 cells/µl, 73% viability) were then processed with 10X Chromium v3.1 and sequenced with DNBSEQ System (MGI) by Genewiz Japan.

#### Single-cell bioinformatic analyses

Raw scRNA-seq reads were mapped onto genome sequences using the CellRanger pipeline (version 6.0.1)^95^. The Drosophila genome and annotation from the Berkeley Drosophila Genome Project, release 6 version 32 (BDGP6.32), were downloaded from the Ensembl Metazoa database^96^. We employed Velocyto (version 0.17.17)^55^ to obtain loom files that describe the spliced/unspliced expression matrices. We merged the loom files with Loompy (version 2.0.16) and converted the merged file into a Seurat object (version 4.0.4)^97^. Quality check and preprocessing were performed using Seurat. We filtered out cells that expressed less than 1,000 or more than 5,000 genes, along with cells with a proportion of mitochondrial RNA larger than 5% from the downstream analysis. We also filtered out hemocytes and visceral muscle cell clusters, as they were considered contamination. Doublets were inferred and removed using DoubletFinder (version 2.0.3)^98^ using standard parameters and the 10X Genomics doublet rate estimation of 0.8%. The remaining 4,184 high-quality cells were normalized and rescaled by regressing on per-cell number of UMIs and mitochondrial content by SCTransform (version 0.3.2)^99^. Dimension reduction was performed by UMAP^100^ using the top-30 principal components from principal component analysis (PCA). We tested multiple resolutions for Louvain graph-based clustering (0.3, 0.5, 0.6, 0.8, 1.0, 1.6), and chose 0.5 for the final fixed resolution. Marker genes were identified using Seurat’s “FindAllMarkers,” with a log fold-change threshold of 0.7 and a minimum percentage of cells of 10%. Gene Ontology term enrichment analysis was performed on the gene sets (p < 0.01, q < 0.01) using ClusterProfiler (version 4.2.2)^101^.

We assessed and annotated the clustering results based on validated markers (**Table S1**). We also compared our annotated clusters to the cell atlas of the adult *Drosophila* midgut^48^ and FACS-sorted EEs^46^ using multidimensional scaling (MDS) scores and combined UMAP coordinates.

Trajectory analysis was performed with scVelo (version 0.2.4)^54^ using “dynamical model” mode, and the UMAP coordinates were imported from the Seurat analysis. The cell fate and terminal state probabilities were calculated considering all clusters using CellRank (version 1.5.1)^56^. For the EE subpopulation identification analysis, we isolated the cluster “AstC^+^EE” and then subjected it to another clustering using the same pipeline with 20 dimensions. Subclustered cell populations AstC^+^EE_0 and AstC^+^EE_1 were renamed on top of AstC^+^EE and used for further comparisons with ISC1, ISC2, and Tk^+^EE clusters.

### SABER FISH

We referred to Kishi et al.^102^ and Amamoto et al.^103^ for probe design, primer concatemerization, and FISH methodology. The probe set for *Dl* was selected from balance type sequences defined in the Oligominer pipeline^104^ (**Table S4**). Concatemerization was performed in the reaction mixture (0.2 U/ml Bst LF polymerase, 2.0 µM primer mix, 0.2 µM Clean G, 1.0 µM hairpin, 0.3 mM dNTPs without dGTP, 10 mM MgSO4) at 37°C for 2 hrs and then at 80°C for 20 min. Concatemers were purified using the MinElute PCR Purification Kit (QIAGEN).

Dissected midguts were fixed with 4% PFA for 30 min at RT, washed with 0.1% Tween-20 at RT, and then with pre-warmed wHyb solution (2×SSC, 1% Tween-20, 40% Formamide) for >15 min at 43°C. Samples were incubated with the primary oligo (1 µg concatemer in 2×SSC, 1% Tween-20, 40% Formamide, 10% Dextran) for 16-24 hrs at 43°C, washed with wHyb at 43°C for 2×30 min, with 2×SSC at 43°C for 2×15 min, then with 0.1% Tween-20 at 37°C for 2×5 min. After incubation with the secondary fluorescent oligo (final 0.2 µM, **Table S4**) at 37°C for 15 min, samples were washed with 0.1% Tween-20 at RT for 2×5 min, then further immunostained at RT for 45 min. Subsequent incubation with secondary antibody was also performed at RT for 45 min. After nuclear staining using Hoechst 33342, samples were mounted in Slowfade Diamond and imaged with confocal microscopy. Following antibodies were used for immunostaining: anti-GFP (MBL, 1:500), anti-dsRed (Clontech 632496, 1:1000). Both antibodies were dissolved in 0.1% Tween-20.

#### Feeding assay

Food intake was measured using cornmeal food containing 1% (w/v) FCF blue dye (Wako, 027-12842). Female adults were fed with the dyed medium for 2 hrs at 18°C and were then homogenized in a 1.5 ml tube containing 150 µl MillQ water (8 flies/tube). Supernatant was collected after centrifugation at 10,000 x g for 10 minutes. Dye content in the supernatant was measured by reading absorbance at 630 nm with Nanodrop 2000c (ThermoFisher). The standard curve was generated by measuring serial dilutions of pure FCF dye (0.00025%, 0.0005%, 0.001%, 0.0025%, 0.005%).

#### *In silico* modeling

The mathematical model predicting each cell number was constructed at the cell population level with continuous variables:

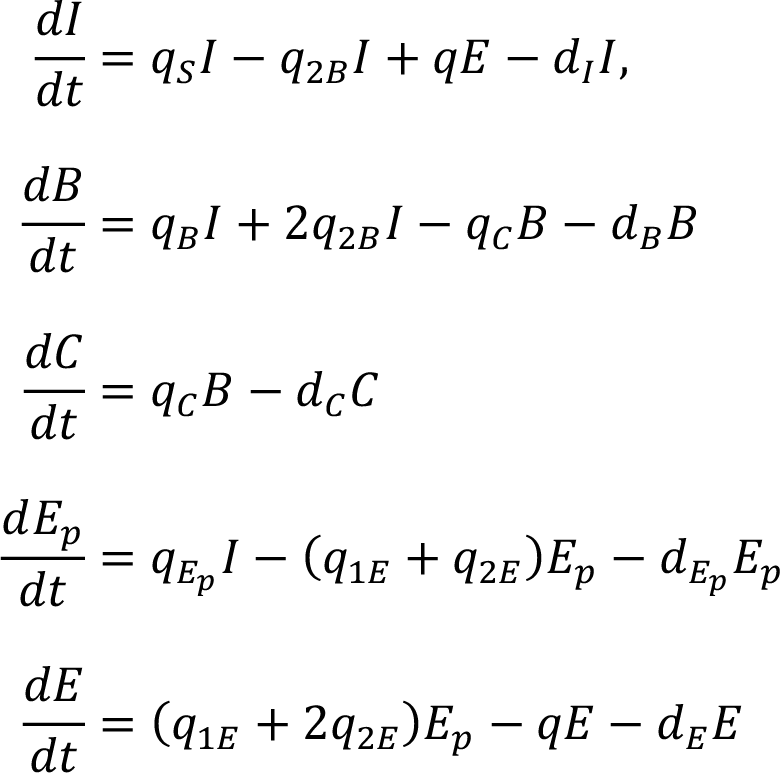

where each term represents cell differentiation and dedifferentiation (**Figure S6A**), and cell death. The variables *I*, *B*, *C*, *E*_*p*_ and *E* represent the number of ISC, EB, EC, EEP and EE cells, respectively, and *t* (day) is time. See **Table S2** for a list of parameter values and see below for definitions of the functions that depend on time *t* or the above variables *I*, *B*, *C*, *E*_*p*_, and *E*.

The cell division rate *a* = *a*(*t*) is defined as:

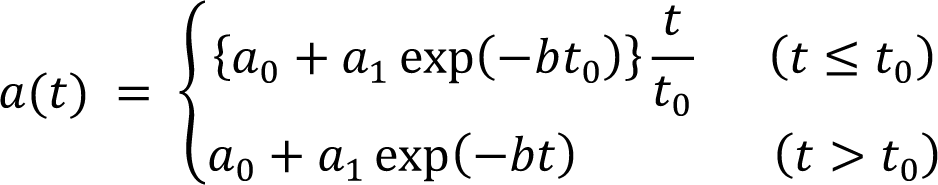

where *a*_<_ is the steady state cell division rate, and the other parameters are estimated from measured mitotic activity (**Figure S6B**). Then the symmetric division rate is *q*_*S*_ = *ap*_*S*_, where *p*_*S*_ is the ratio of symmetric division. The asymmetric division rate *q*_*B*_ = *ap*_*B*_, *q*_*E*__*p*_ = *ap*_*E*__*p*_ and the symmetric differentiation rate *q*_*2B*_ = *ap*_*2B*_ are defined similarly. Note that *p*_*S*_ + *p*_*B*_ + *p*_*E*__*p*_ + *p*_*2B*_= 1. Each division ratio *p*_*i*_ varies piecewise linearly in time (**Figure S6C**):

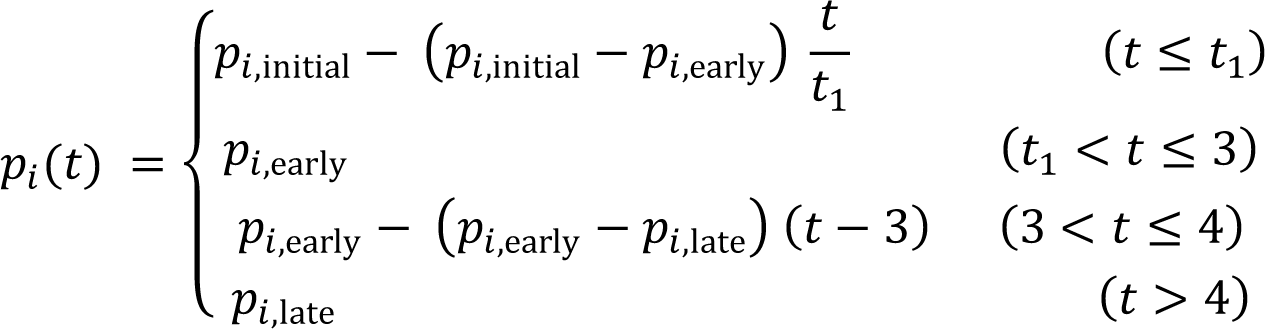

where *i* = S, *B*, *E*_*p*_, 2*B*.

The rate of dedifferentiation *q* reaches a maximum value at day 1, then decays, and is zero after day 4 (**Figure S6D**):

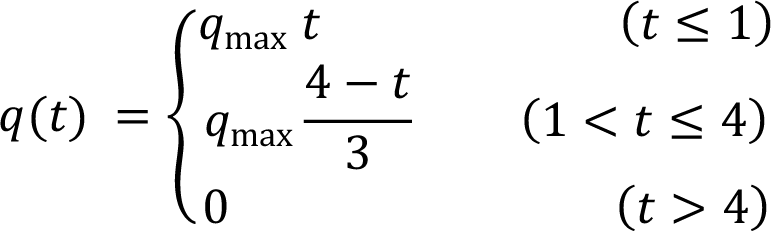

The differentiation rate *q_C_* from EBs to ECs also reaches its maximum at day 1 and then decreases over time. Conversely, the cell death rate *d_B_* of EBs increases over time^105^. The time changes after day 1 are described by the Hill function (**Figure S6E**):

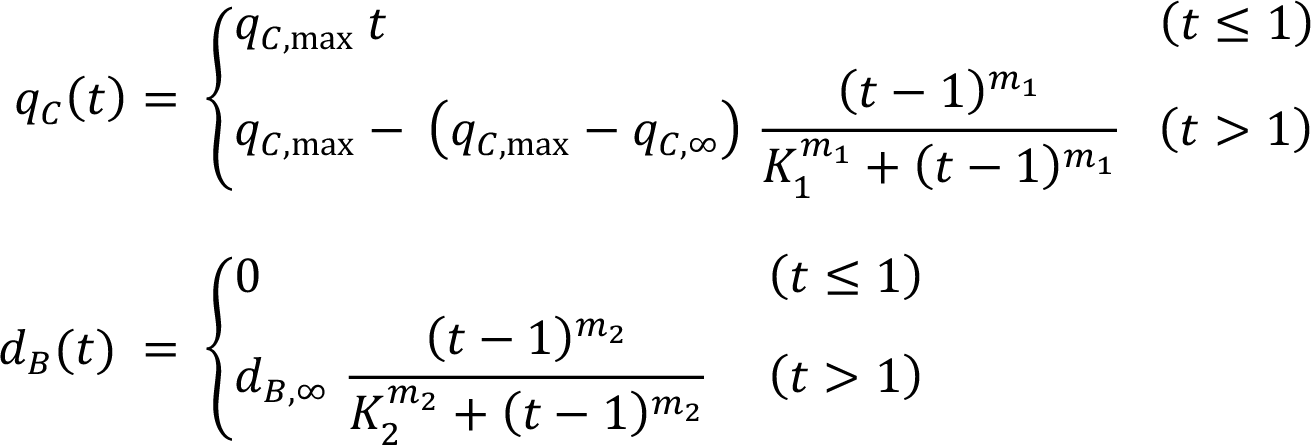

The rate constants *q_C,∞_*, *d_B,∞_*, and the cell death rates *d*_1_, *d_C_*, *d_E_* are determined by steady state conditions:

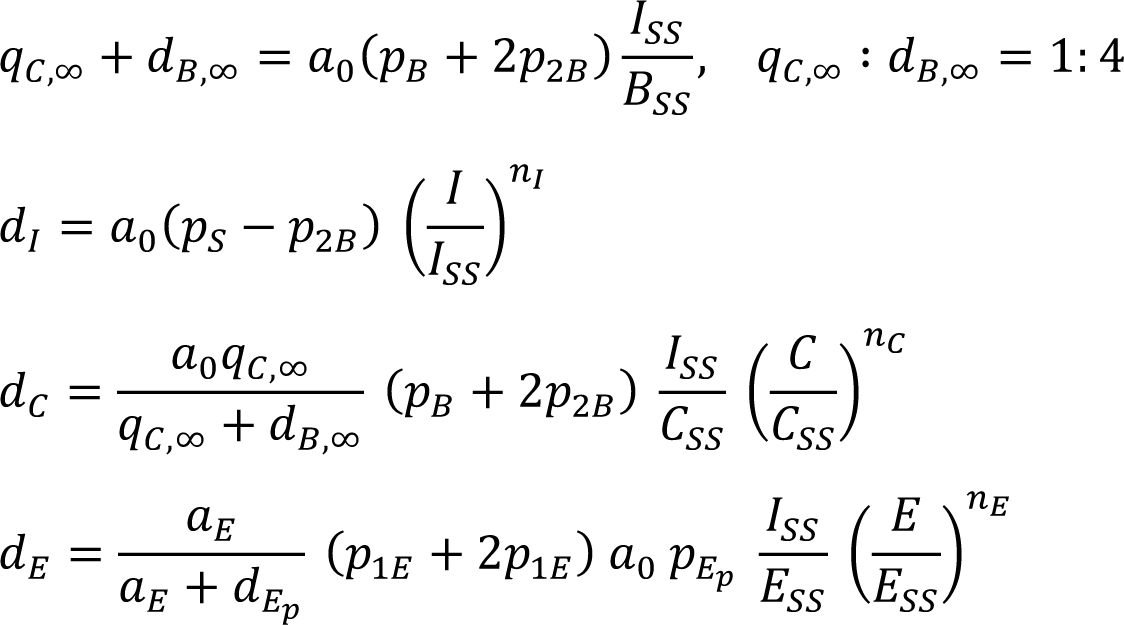

where *I_SS_*, *B_SS_*, *C_SS_*, and *E_SS_* represent the steady state values of *I*, *B*, *C*, and *E*, respectively, and are determined by (Marianes and Spradling, 2013)^11^:

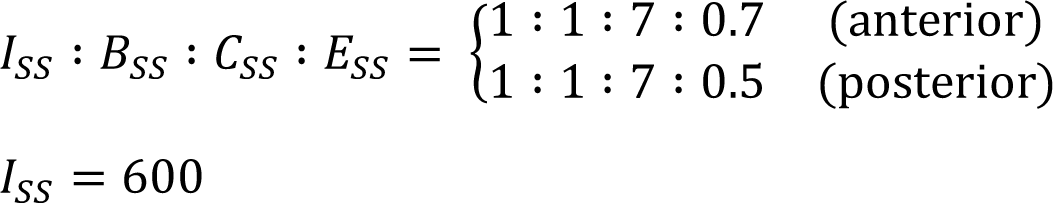

#### RT-qPCR

Total RNA was purified from 10-15 midguts using the ReliaPrep RNA Tissue Miniprep System (Promega). cDNA was made from 100 or 200 ng of RNA using PrimeScript RT Reagent Kit (TaKaRa). Quantitative PCR was performed using TB Green Premix Ex Taq II (TaKaRa) and a QuantStudio 6 Flex Real-Time PCR System (ThermoFisher). *RpL32* was used as an internal control. Primer sequences were listed in **Table S4**.

### QUANTIFICATION AND STATISTICAL ANALYSIS

#### Boundary between midgut compartments

The midgut region (anterior, middle, and posterior) was determined based on defined morphological characteristics^9, 11^. We first searched for characteristic constrictions at the boundary between the anterior-middle and middle-posterior. We also verified these boundaries by checking the length of each region (the ratio of length, anterior:middle:posterior, is roughly 4:1:4). We focused on the anterior and the posterior midgut given the different lineage hierarchy in the middle midgut^28, 29^.

#### Twin spot clone type

In twin spot MARCM experiments (**Figure 1D, 1E, and S1H**), heat shock induces mitotic recombination that results in clonal labeling of one ISC daughter with GFP and the other daughter with RFP. Both fluorescent proteins are expressed by ubiquitous promoter, thus visualizing clonal expansion of the two ISC daughters individually^3, 31, 33^. Symmetric ISC division generates two ISCs that undergo additional rounds of mitosis. We therefore classified symmetric division as when both the GFP clone and RFP clone contain ≥ 2 cells (total ≥ 4 cells in a twin spot). On the other hand, asymmetric ISC division generates one ISC and one differentiated cell that loses mitotic activity. We therefore classified asymmetric division as when either color consists of only one cell and the other color contains ≥ 2 cells (total ≥ 3 cells in a twin spot). We excluded twin spots with only one cell in both colors (total 2 cells in a twin spot) from the quantification, since we cannot distinguish whether the singly labeled cell is a differentiating cell or an ISC that does not undergo additional mitosis. We also excluded single-color clones without an adjacent clone of the opposite color (e.g., GFP clone without adjacent RFP clone, Figure S1I), which likely arise from cell death in one color.

Although a subset of rare EEPs also exhibit mitotic activity in addition to ISCs^38^, EEPs can divide only once, and resultant daughters are post-mitotic EEs. Thus, if mitotic recombination occurs in EEPs, both colors remain a one cell clone (total 2 cells in a twin spot). We excluded 2-cell twin spots as described above, thus focusing on twin spots originated from ISC division.

#### Quantification of cellular shape

Cell shape (**Figure 3A, 3B, S3I**) was quantified using Fiji software. The cell membrane was visualized by anti-Armadillo staining and recorded as the ROI with the polygon selection tool. The circularity of ROIs was measured using the Shape descriptors plugin. High circularity indicates a rounded shape (similar to a complete circle) whereas low circularity indicates an angular and/or elongated shape. Cell type was determined by combining anti-Pros staining, *esg-lacZ* reporter, and lineage tracing using *pros-Gal4*: EEs were Pros^+^β-gal^−^, esg^+^ cells were Pros^−^β-gal^+^lineage^−^, and EE-derived esg^+^ cells were Pros^−^β-gal^+^lineage^+^.

#### Quantification of Dl^+^ cell ratio

The Dl^+^ cell ratio (**Figure 5E, S5E, and S5F**) was measured by counting the total cell number as well as the Dl^+^ cell number using Fiji. Quantification of total cell number was performed as follows: (1) Remove noise signal of Hoechst staining with the Despeckle command. (2) Binarize using the Threshold command. (3) Fill stainless nuclear compartments such as the nucleolus using the Fill Holes command. (4) Divide multiple nuclei that are continuously adjacent using the Watershed command. (5) Measure the number of nuclei using the Analyze Particles command. The Dl^+^ cells were defined as diploid cells with membrane or punctate Dl signal.

#### Quantification of midgut size

The midgut area (**Figure 5G, S5G-S5J, and S5P**) was measured using a previously established macro for Fiji^77^. Briefly, staining artifacts and fluorescent signal of other tissues (Malpighian tubules and trachea) were cut out using the line tool. Then the midgut ROI was selected and binarized. The size, length, and thickness of selected ROIs were measured automatically.

#### Statistics

Statistical analyses were performed using Excel and RStudio. Two tailed *t* tests were used for comparisons between two groups. One-way ANOVAs with post hoc Tukey tests were performed when comparing three or more groups. chi-square tests were used for comparisons for the symmetric-asymmetric ratio (**Figure 1E**) and the ratio of EC-only clones (**Figure 7H**). Significance is indicated in the figures as follows: *P≤0.05, **P≤0.01, ***P≤0.001, Not Significant (N.S.): P>0.05. Bar graphs show mean ± standard error. Boxplots show median (thick line in the box), first and third quartiles (bottom and top of the box), minimum value (lower whisker), and maximum value (upper whisker). Dots in bar graphs and boxplots indicate individual values. Violin plots indicate distribution of individual values.

### Supplemental Tables

**Table S1.** Marker genes utilized for cell type annotation.

| Gene symbol | Cell type |
| --- | --- |
| Dl | ISC |
| esg | ISC/EB |
| Su(H) | EB |
| pros | EE |
| Tk | Tk<sup>+</sup>EE |
| NPF | Tk<sup>+</sup>EE |
| DH31 | Tk<sup>+</sup>EE |
| AstC | AstC<sup>+</sup>EE |
| AstA | AstC<sup>+</sup>EE |
| CCHa1 | AstC<sup>+</sup>EE |
| Orcokinin | AstC<sup>+</sup>EE |
| alphaTry | Anterior EC (aEC) |
| betaTry | Anterior EC (aEC) |
| LambdaTry | Posterior EC (pEC) |
| iotaTry | Posterior EC (pEC) |
| Vha100-4 | Middle EC (mEC) |
| Hml | Hemocyte |
| zfh1 | Hemocyte |
| vkg | Visceral muscle |
| Mhc | Visceral muscle |
| Mlc2 | Visceral muscle |

**Table S2.**
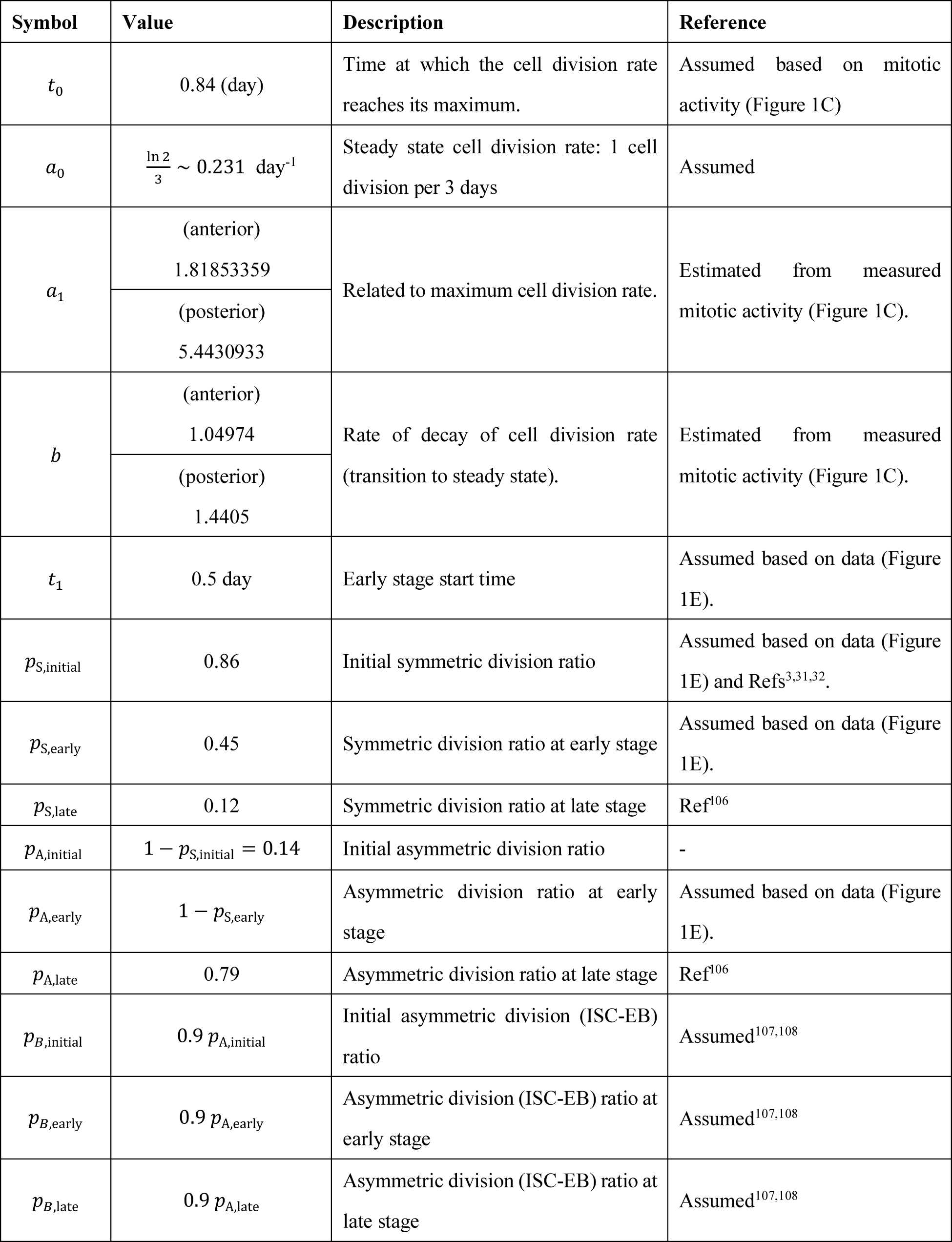
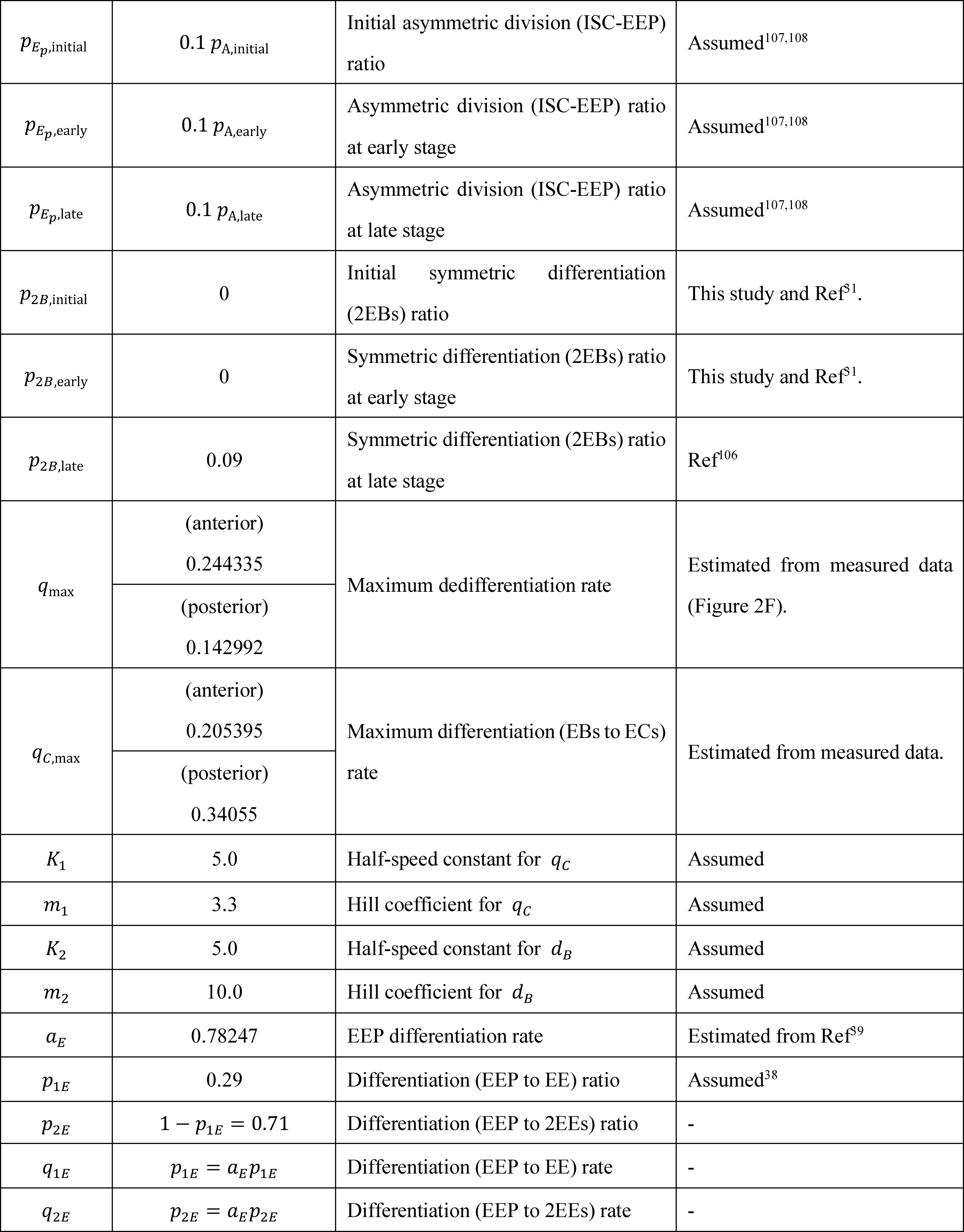
List of parameters used in the simulation.

**Table S3.**
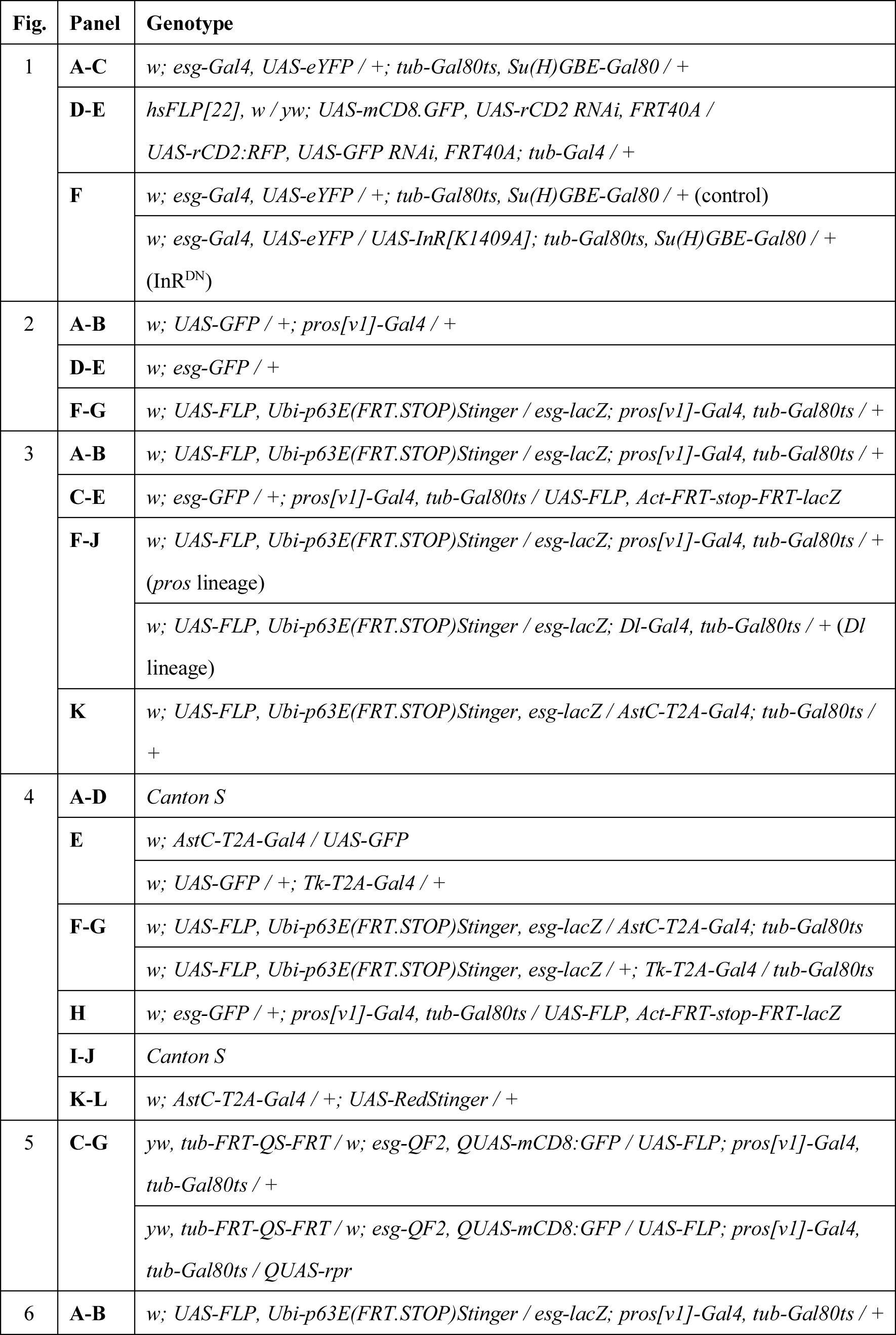
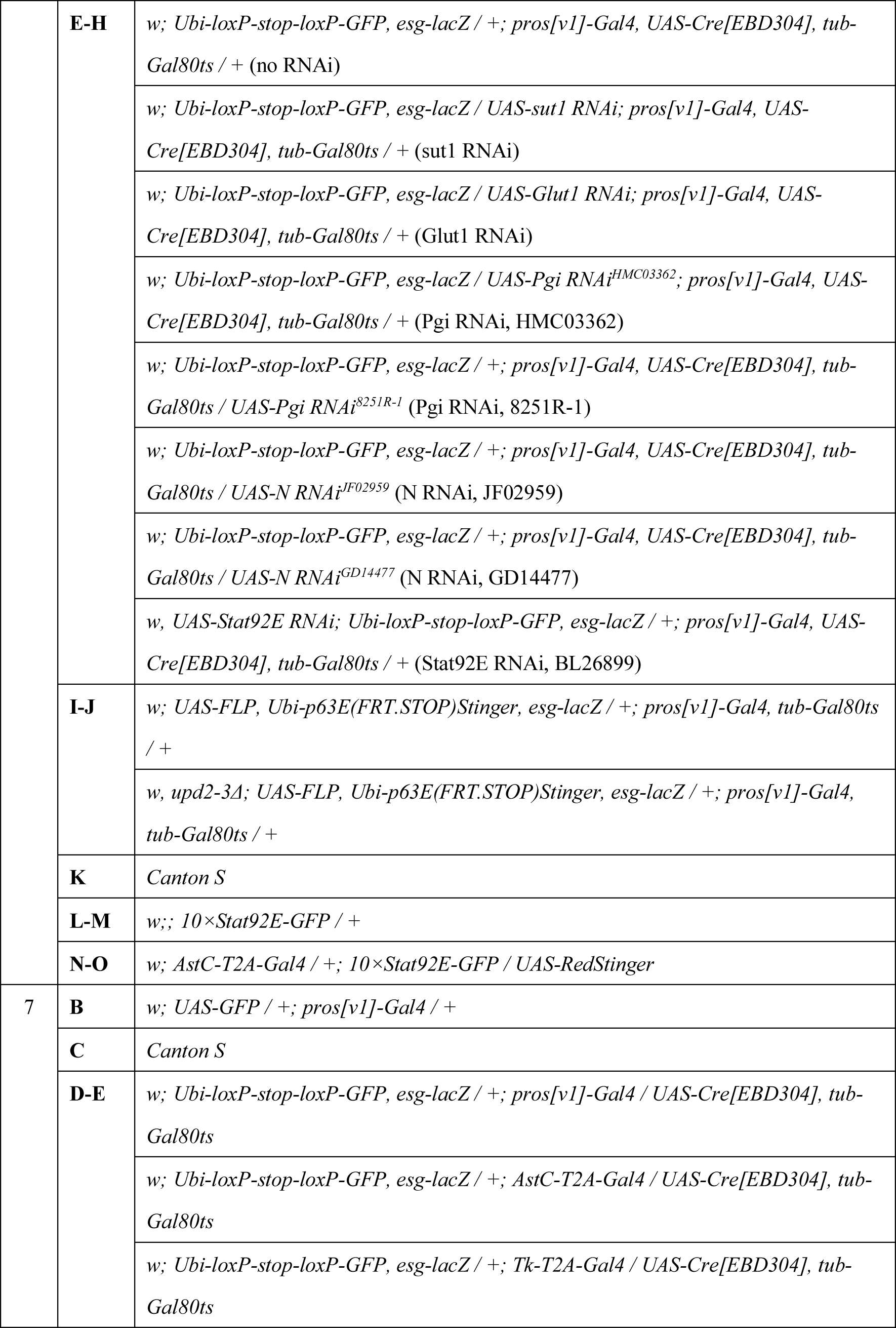
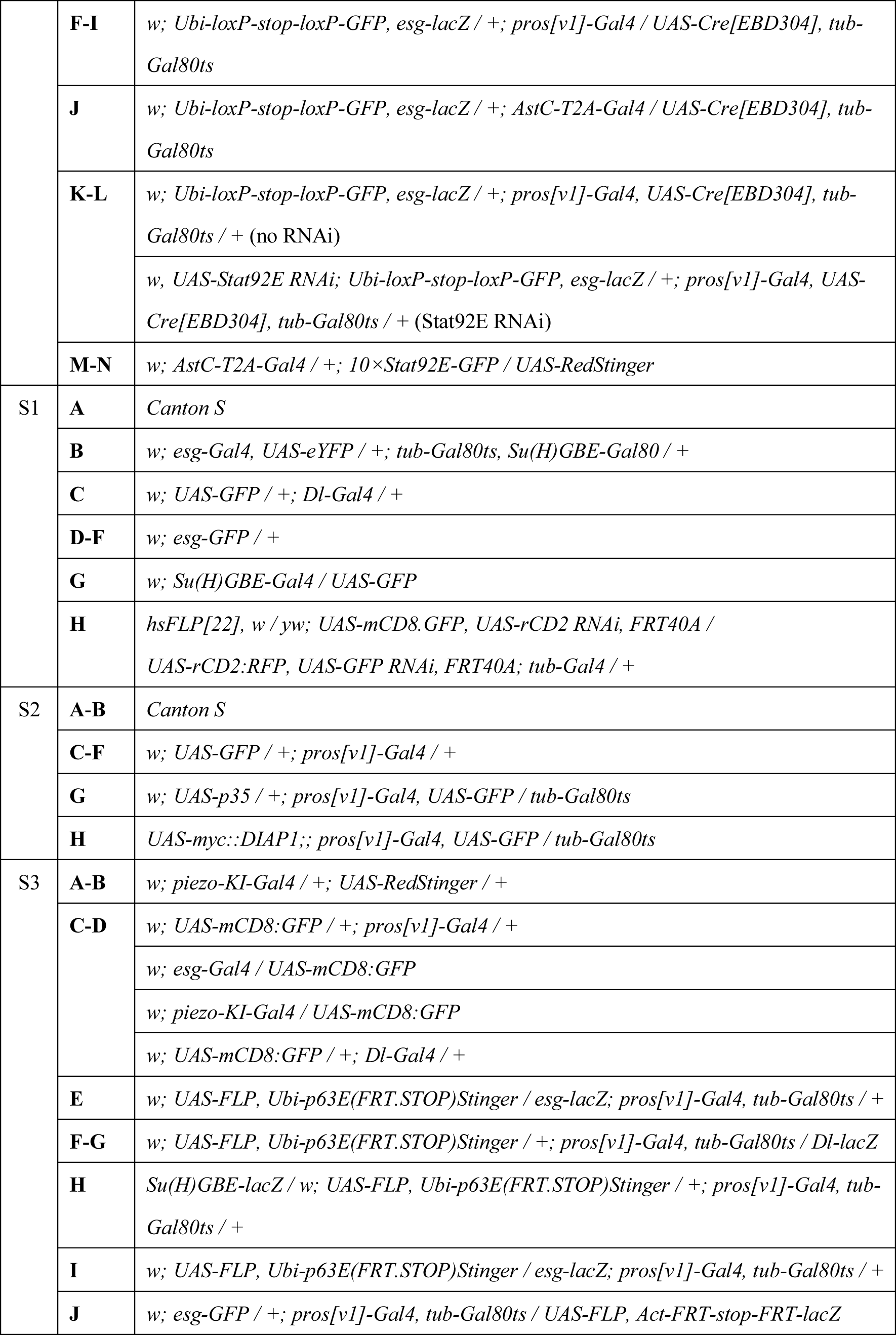
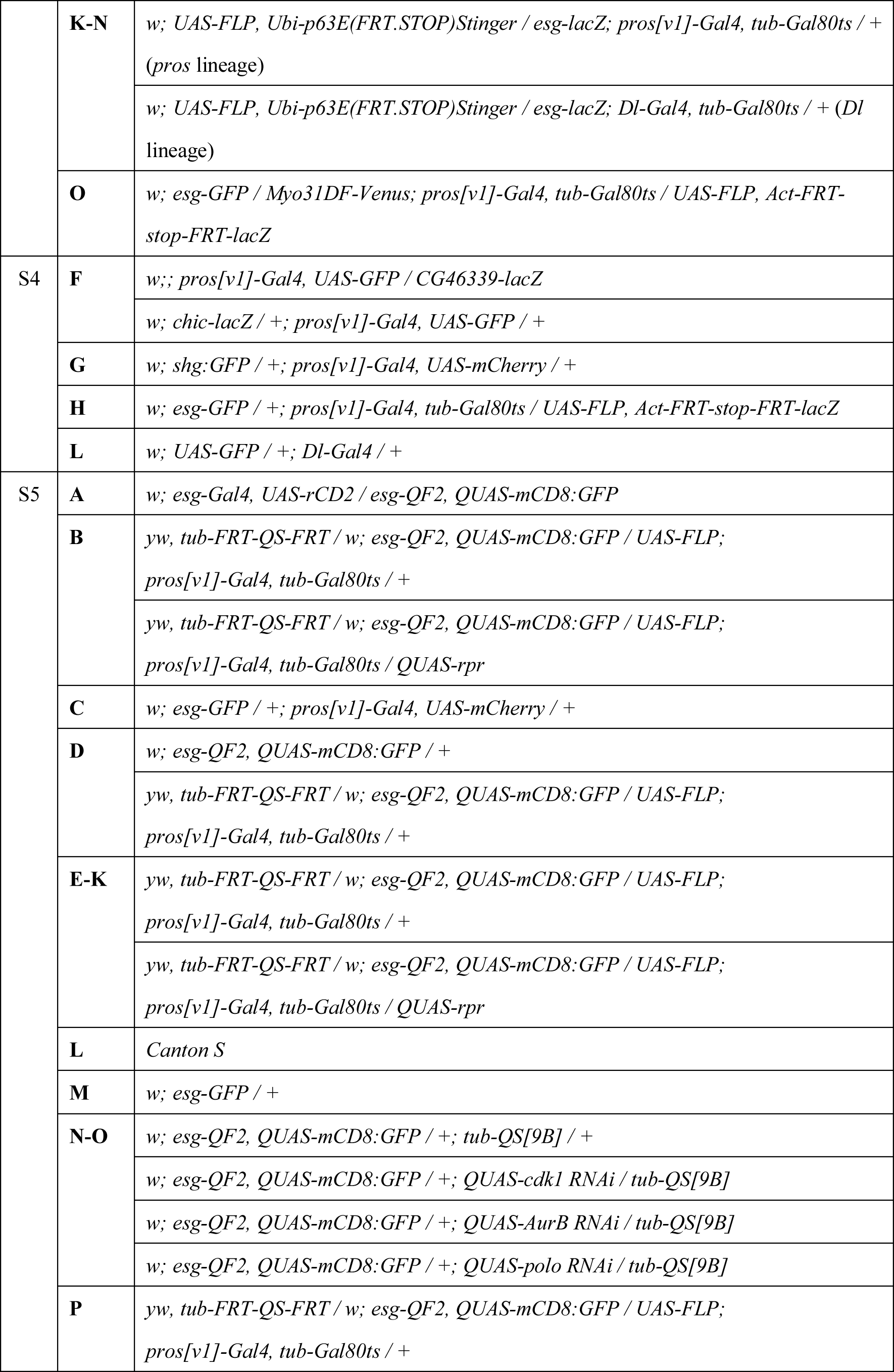
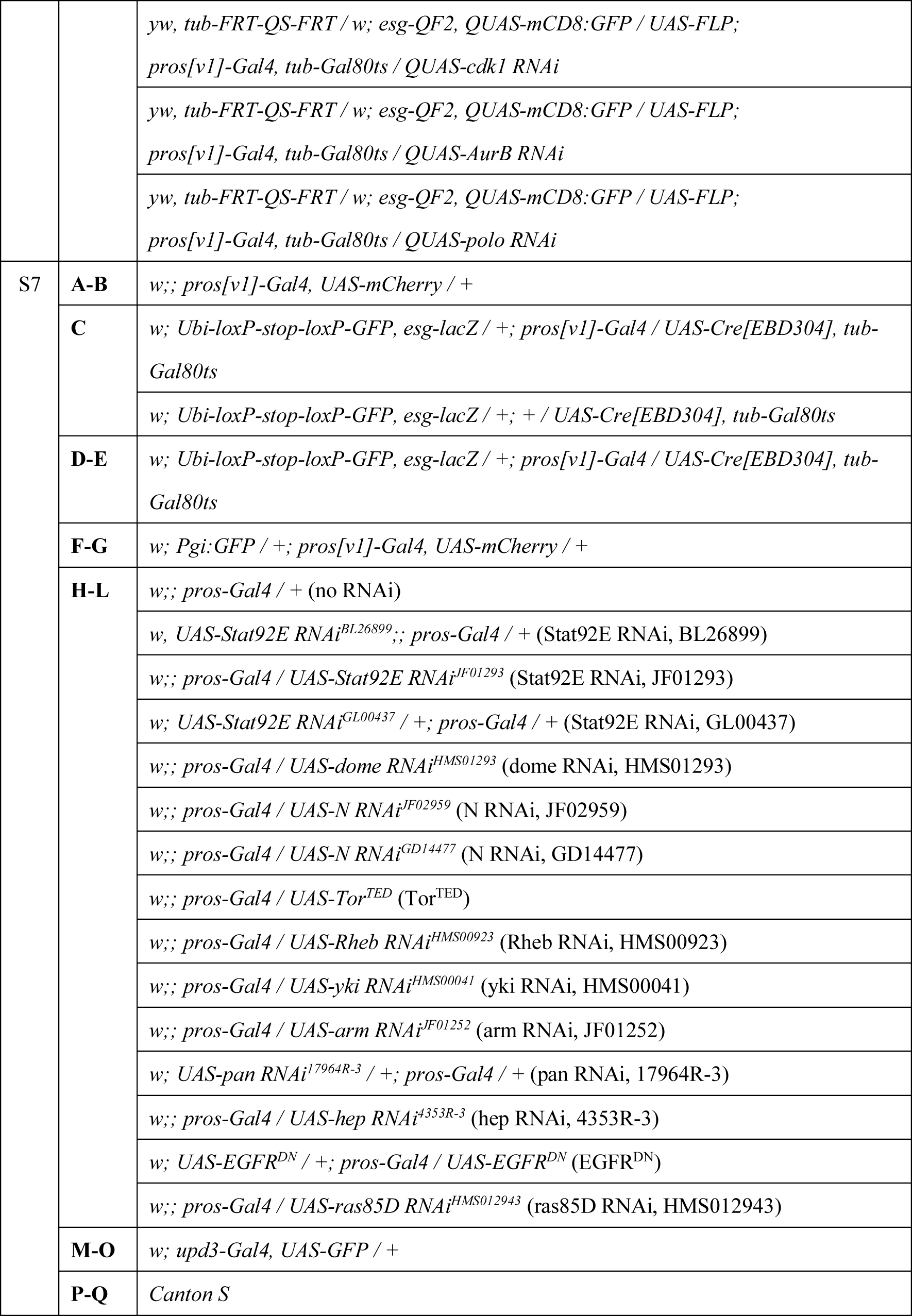
Detailed genotypes in each experiment.

Table S4. Oligo sequences

Table S5. Absolute cell counts for main figures

Table S6. Absolute cell counts for supplemental figures

*Table S1-S3 are included in this file, and Table S4-S6 are separately uploaded as Excel spreadsheets.

## Supplemental figure legends

**Figure S1. The number of ISCs and EBs increases after eclosion.**

(A) Total cell number in the anterior midgut. In the fed condition, the total cell number increased in a feeding dependent manner between Day 1 and Day 3. In the starved condition, the total cell number increased between Day 0 and Day 1; however, there was no further increase between Day 1 and Day 3. Fed: n=10 (Day 0), 9 (Day 1), 11 (Day 2), 12 (Day 3), 12 (Day 7). Starved: n=11 (Day 0), 9 (Day 1), 11 (Day 3).

(B) The absolute number of *esg^+^Su(H)^−^* cells in Day 1, Day 2, and Day 3 guts (related to Figure 1B). n=13 (Day 1), 11 (Day 2), 10 (Day 3) midguts.

(C) The number of *Dl-Gal4>GFP^+^* cells and the mitotic activity of *Dl>GFP^+^* cells. The number of *Dl>GFP^+^* cells similarly increases both in the anterior/posterior midgut, however, their mitotic activity is lower in the anterior midgut than in the posterior midgut. n= 12 (Day 1), 11 (Day 2), 13 (Day 3) midguts.

(D) The absolute number of *esg-GFP^+^*ISCs/EBs in Day 1, Day 2, and Day 3 guts. n=9 (Day 1), 12 (Day 2), 10 (Day 3) midguts.

(E) The relative number and the mitotic activity of *esg-GFP^+^* ISCs/EBs. While the number of ISCs/EBs increases ∼1.5 fold both in anterior and posterior midguts, the mitotic activity of *esg-GFP^+^* cells is significantly lower in the anterior midgut than in the posterior midgut.

(F) There is no increase in ISC/EB number under starved condition. n=14 (Day 1), 10 (Day 2), 12 (Day 3) midguts.

(G) The number of *Su(H)GBE-Gal4>GFP^+^* EBs increases after eclosion in both midgut regions in the fed condition. n=11 (Day 1), 11 (Day 2), 10 (Day 3) midguts.

(H) Representative image of a non-twin clone (white arrows) that exhibits only one fluorescence type in the twin-spot MARCM system. The typical twin-color clone is indicated by yellow arrows. The right graph shows quantification for the ratio of the non-twin clones in all clones. Scale bar: 50 µm.

Not Significant (N.S.): P>0.05, **P≤0.01, ***P<0.001. One-way ANOVAs with post hoc Tukey tests.

**Figure S2. The feeding-dependent and apoptosis-independent decline in EE number in the early adult midgut.**

(A and B) The number of EEs is measured by anti-Prospero staining. Prospero^+^ cells decrease in the fed condition (A) but not in the starved condition (B). n=10 (Day 0), 9 (Day 1), 11 (Day 2), 12 (Day 3), 12 (Day 7) midguts in (A), and n=11 (Day 0), 9 (Day 1), 11 (Day 2), 11 (Day 3) midguts in (B).

(C) Representative images of TUNEL staining. Paraquat (PQ) feeding acts as a positive control for midgut cell death. *pros>GFP^+^*cells rarely exhibit TUNEL signal. Scale bar: 100 µm.

(D and E) Quantification of TUNEL signal. PQ feeding significantly increases the number of TUNEL^+^ cells, suggesting that TUNEL staining successfully detects apoptotic events (D). TUNEL^+^ EEs are rare both in PQ treated guts and early adult guts (E). n=6 (PQ), 8 (Day 1), 6 (Day 2) midguts.

(F) Sytox staining, which detects the membrane permeability characteristic of dead cells, is rarely detected in EEs. Paraquat feeding acts as a positive control for midgut cell death. Scale bar: 20 µm.

(G) Overexpression of *p35* does not inhibit the decrease of EE number after eclosion. n=11 (Day 1), 10 (Day 2), 6 (Day 3) midguts.

(H) Overexpression of *Diap1* does not inhibit the decrease of EE number after eclosion. n=8 (Day 1), 5 (Day 2), 15 (Day 3) midguts.

N.S., not significant: P>0.05, *P≤0.05, **P≤0.01, ***P≤0.001. One-way ANOVAs with post hoc Tukey tests.

**Figure S3. Direct conversion from mature EEs into ISCs.**

(A) The Pros^+^*piezo^+^* EEPs are detected in midguts 3 days after puparium formation (APF), but not in those 4 days APF. The *piezo-KI-Gal4>RFP* pattern reproduces the data in previous report^40^.

(B) Quantification of Pros^+^*piezo^+^*cells among Pros^+^ cells. Pros^+^*piezo^+^* EEPs are rarely detected in midguts 4 days APF. n=15 (3 days), 28 (4 days) images.

(C) Representative images of apical protrusion in mature EEs. The morphology of Pros^+^ cells were examined by expressing mCD8:GFP with *Gal4* drivers that mark pan-EE lineage (*pros-Gal4*) or immature EE progenitors (*esg-Gal4*^40, 61^, *Dl-Gal4*^38, 39^, *Piezo-KI- Gal4*^40^) to see the apical protrusion, which was proposed as a characteristic of differentiated EEs^43, 82^. In the adult midguts, Pros^+^ cells that are labeled by *pros-Gal4* extend cellular protrusion toward the apical lumen, while those marked with *esg-Gal4, Dl-Gal4*, or *Piezo-KI-Gal4* lack this structure and exhibit round shape. At 4d APF, Pros^+^ cells that are marked with *pros-Gal4* also exhibit the apical protrusion, suggesting that Pros^+^ cells complete maturation into EEs before eclosion.

(D) The length of apical protrusion was quantified by using z-stack images of Pros^+^ cells. We measured the length from the apical tip of nuclear Hoechst signal to the apical tip of mCD8:GFP signal.

(E) Whole midgut image of pros-lineage tracing sample (genotype: *pros-Gal4, tub- Gal80ts>UAS-FLP, Ubi-FRT-stop-FRT-GFP*). No leaky labeling is detected at Day 7 when flies were kept at 18°C, while temperature shift to 29°C (Figure 2C) induces GFP^+^ cells.

(F) A subset of *pros-*lineage cells loses Pros expression and instead acquires *Dl* expression after eclosion (arrowhead). Experimental scheme is the same as in Figure 2C.

(G) Quantification of *Dl^+^* ratio in *pros*-lineage cells in fed samples. n=6 (Day 1), 11 (Day 4) midguts.

(H) *pros*-lineage cells rarely exhibit *Su(H)* expression in Day 1 fed guts and Day 4 fed guts. n=6 (Day 1), 4 (Day 4) midguts.

(I) Quantification of Figure 3A for posterior midgut. Circularity of EEs, EE-derived esg^+^ cells, and non-EE-derived esg^+^ cells were quantified. n=21 (Day 1, EE), 9 (Day 1, EE- derived esg^+^), 13 (Day 1, non-EE-derived esg^+^), 23 (Day 4, EE), 12 (Day 4, EE-derived esg^+^), 10 (Day 4, non-EE-derived esg^+^) cells.

(J) Quantification of Figure 3C. CCHa1 intensity is significantly higher in EE-derived (lineage^+^) esg^+^ cells compared to non-EE-derived (lineage^−^) esg^+^ cells in the Day 1 anterior midgut. n=14 (Day 1, lineage^−^esg^+^), 18 (Day 1, lineage^+^esg^−^CCHa1^+^), 12 (Day 1, lineage^+^esg^+^), 5 (Day 4, lineage^−^esg^+^), 8 (Day 4, lineage^+^esg^−^CCHa1^+^), 6 (Day 4, lineage^+^esg^+^) cells.

(K) The number of cells per clone at Day 1, Day 4, and Day 7.

(L-N) The ratio of esg^+^ cells (L), esg^−^ polyploid cells (M), and esg^−^ diploid cells (N) in the Day 7 *pros-*lineage clones (EE-derived ISCs) and *Dl-*lineage clones (resident ISCs). n=32 (EE-derived ISC, anterior), 42 (resident ISC, anterior), 34 (EE-derived ISC, posterior), 25 (EE-derived ISC, posterior) clones.

(O) Myo31DF-Venus (Myo1A-Venus) localizes to the apical membrane in the *pros-* lineage polyploid cell (arrow). The *pros-*lineage esg^+^ cell (arrowhead) is also detected adjacent to the *pros-*lineage polyploid cell. The subcellular localization of Myo31DF-Venus is similar to that of anti-Myo1A and Myo1A^CPTI004107^ protein trap line^45, 81^.

N.S., not significant: P>0.05, *P≤0.05, ***P≤0.001, One-way ANOVAs with post hoc Tukey tests. Scale bar: 50 µm (A), 500 µm (E), 10 µm (F), and 25 µm (O).

**Figure S4. Validation of clusters annotations and gene signature in EEs.**

(A) Integrated UMAP plot of our single cell dataset with that of Hung et al^48^. Datasets were normalized by SCTransform before Louvain clustering. Clusters in our dataset are shown with bright colors while those in Hung et al. are shown in gray.

(B) MDS plot, together with the UMAP plot, indicates the correlation between our clusters and those of Hung et al.

(C) Neuropeptide expression pattern in our dataset. Our AstC^+^EEs highly express neuropeptides of class I EE (AstC, AstA, Orcokinin, CCHa1, CCHa2)^46^. Similarly, Tk^+^EEs in our dataset express neuropeptides of class II EE (Tk, NPF, Dh31)^46^. Neuropeptides of class III EE (sNPF, CCHa2) are expressed in our AstC^+^EEs, suggesting that class III EEs are not separated in our dataset.

(D) Gene ontology enrichment for ISC1 over ISC2.

(E) Gene ontology enrichment for AstC^+^EEs over Tk^+^EEs. *CG46339, chic, Shg,* and *His2Av* are included in the term “somatic stem cell population maintenance.”

(F) Differential expression of *CG46339* and *chic* is detected in EE population in Day 1 fed guts. Enhancer trap lines *CG46339-lacZ* and *chic-lacZ* were used.

(G) Differential expression of Shg (Drosophila E-Cadherin) is detected in EE population in Day 1 fed guts. Protein trap line Shg:GFP was used. Note that *pros>mCherry^+^* EEs exhibit a round shape, which is consistent with the observation by anti-Armadillo staining (Figure 3A). No obvious differences in His2Av expression were detected *in vivo*.

(H) Quantification of Figure 4H. Intensity of CCHa2, but not of Tk or NPF, is significantly high in EE-derived (lineage^+^) esg^+^ cells compared to non-EE-derived (lineage^−^) esg^+^ cells in Day 1 anterior midgut. CCHa2: n=10 (lineage^−^esg^+^), 12 (lineage^+^esg^−^CCHa2^+^), 7 (lineage^+^esg^+^) cells. Tk: n=12 (lineage^−^esg^+^), 8 (lineage^+^esg^−^Tk^+^), 18 (lineage^+^esg^+^) cells. NPF: n=13 (lineage^−^esg^+^), 22 (lineage^+^esg^−^NPF^+^), 18 (lineage^+^esg^+^) cells.

(I) Subclustering of AstC^+^EE using the same approach for the initial cells clearly reflects the presence of two subpopulations with distinct features.

(J, K) Integrated UMAP plot of our single cell dataset with that from Guo et al^46^. All of our quality-filtered cells (J) and only EEs and ISC1 (K) are merged with FACS-sorted EEs^46^ (Guo et al., 2019). Datasets were normalized by SCTransform before Louvain clustering. Clusters in our dataset are shown with bright colors while those in Guo et al. are shown in gray.

(L) Validation of *Dl* probe set. *Dl* mRNA signal is detected in *Dl-Gal4>GFP^+^* cells.

(M) Expression levels of *CG46339*, *chic*, and *shg* in AstC^+^EE subpopulations, Tk^+^EE, and ISCs. Expression of *CG46339* gradually decreases along AstC^+^EE_1, AstC^+^EE_0, and ISCs compared with the acute down-regulation between ISCs and Tk^+^EE. *chic* and *shg* are upregulated in AstC^+^EE_0 and ISC1 over AstC^+^EE_1.

(N) Expression levels of *dome, Stat92E*, and *socs36E* in AstC^+^EE subpopulations, Tk^+^EE, and ISCs. The dedifferentiating AstC+EE_0 highly expresses genes related to the JAK-STAT pathway compared to Tk+EE.

N.S., not significant: P>0.05, *P≤0.05. One-way ANOVAs with post hoc Tukey tests. Scale bar: 5 µm.

**Figure S5. Validation of the ablation system and growth defect by mitotic inhibition in EE-derived ISCs.**

(A) The newly established *esg-QF2* recapitulates its original *esg-Gal4* pattern. Arrows: *QF2^+^Gal4^−^* cells, arrowhead: *QF2^−^Gal4^+^* cell. n=11 (anterior), 12 (posterior) images.

(B) EE-derived *esg^+^* cells are detected in 4-day fed guts (upper and lower left panels) and are eliminated by *rpr* overexpression (middle and lower right panels). These GFP-marked cells are diploid, a characteristic of *esg^+^* ISCs. Scale bars: 500 µm (upper and middle panels), 50 µm (lower panels).

(C) *pros* is highly expressed in adult brain cells whereas esg^+^ cells are rare.

(D) *esg-QF2>mCD8:GFP* signal is absent in most brain cells, except for a few cells in the subesophageal ganglion (upper panels). *pros*-derived *esg^+^* cells are completely absent in the adult brain (lower panels).

(E) Ablation effect on Dl^+^ cell ratio depends on nutrient intake after eclosion. *G: GFP* (control), *Gr: GFP+rpr* (ablation), n=22 (*G*), 22 (*Gr*) images analyzed.

(F) Ablation effect on Dl^+^ cell ratio depends on the priming of *rpr* overexpression. n=28 (*G*), 26 (*Gr*) images analyzed.

(G) Ablation effect on midgut size depends on nutrient intake after eclosion. n=15 (*G*), 13 (G*r*) midguts.

(H) Ablation effect on midgut size depends on the priming of *rpr* overexpression. n=15 (*G*), 15 (*Gr*) midguts.

(I, J) Ablation of EE-derived esg+ cells impaired the midgut growth in thickness, but not in length. n=15 (*GFP,* Day 1), 15 (*GFP*, Day 10), 12 (*GFPrpr*, Day 1), 12(*GFPrpr*, Day 10) midguts.

(K) Food intake in 2 hours was measured at Day 1, Day 4, and Day 10 after eclosion. *rpr* induction did not decrease the amount of blue dye ingestion. *G: GFP* (control), *Gr: GFP+rpr* (ablation). n=8 (Day 1), 9 (Day 4), 7 (Day 10) experiments. Eight flies were used for each sample.

(L) Feeding assay detects decreases in food intake. Wild type adults consumed blue dye food for 20 minutes or 2 hours. Food intake in 20 minutes is significantly less than that in 2 hours. n=9 experiments. Eight flies were used for each sample.

(M) Pros^+^esg^+^ EEs in the middle midgut do not exhibit PH3 signal (0/24 PH3^+^ cells from 11 midguts). Arrows: Pros^+^esg^+^ EEs, arrowheads: PH3^+^ cells.

(N) Mitotic inhibition using *esg-QF2, tub-QS* system and the newly established QUAS- RNAi lines targeting *cdk1, AurB*, and *polo*. Mitotic inhibition causes mis-differentiation of *esg-QF2>GFP^+^* cells in the anterior midgut, but not in the middle midgut. Adult flies were fed with quinic acid for 7 days before experiments.

(O) Quantification for (N). The mis-differentiation phenotype (e.g., abnormal endoreplication) is quantified by nuclear size. Anterior: n=140 (control), 144 (*cdk1 KD*), 68 (*AurB KD*), 65 (*polo KD*) cells. Middle: n=94 (control), 95 (*cdk1 KD*), 205 (*AurB KD*), 132 (*polo KD*) cells. Posterior: n=113 (control), 160 (*cdk1 KD*), 149 (*AurB KD*), 232 (*polo KD*) cells.

(P) Mitotic inhibition in EE-derived esg^+^ cells impairs growth of the anterior midgut. No significant effect is exhibited in the posterior midgut. n=18 (control, Day 1), 21 (control, Day 10), 17 (*cdk1 KD*, Day 1), 20 (*cdk1 KD*, Day 10), 16 (*AurB KD*, Day 1), 18 (*AurB KD*, Day 10), 19 (*polo KD*, Day 1), 24 (*polo KD*, Day 10) guts.

N.S., not significant: P>0.05, **P≤0.01, ***P≤0.001. One-way ANOVAs with post hoc Tukey tests (E-H), two tailed *t* test (I-L). Scale bars: 50 µm (A, N), 200 µm (C-D), 20 µm (M).

**Figure S6. Mathematical model of cell population dynamics in the adult midgut.**

(A) Pathways of cell differentiation and dedifferentiation in the mathematical model.

(B) Cell division rate *a*, fitted with mitotic activity data (Figure 1C).

(C) Symmetric division ratio *p*_*S*_ in the anterior region, based on measured data (Figure 1E) and previously reported data^3, 31, 32^. Other parameters denoted in the form of *p*_*i*_ are defined by similar piecewise linear functions.

(D) Dedifferentiation rate *q*. Its maximum was assumed to be taken at exactly Day 1 and was estimated from 0-1-day data (Figure 2G).

(E) EB to EC differentiation rate *q_C_* and EB death rate *d_B_* . The maximum differentiation rate *q*_1,max_ was assumed to be taken at exactly Day 1 and was estimated from data.

**Figure S7. Glucose incorporation and the JAK-STAT pathway underlie EE dedifferentiation.**

(A and B) Anterior EEs incorporate more 2-NBDG than do posterior EEs, which is quantified in (B). 2-NBDG is orally treated between Day 0 to Day 1. n=68 (anterior), 44 (posterior) *pros>mCherry^+^* cells.

(C) No leaky labeling is detected in T-trace midguts at Day 14. Experimental scheme indicated in Figure 7A is applied. Detailed genotype of *pros^ts^>T-trace* is *pros-Gal4, tub- Gal80ts, UAS-Cre^EBD^, Ubi-loxP-stop-loxP-GFP*. In “pros^ts^>T-trace, no estrogen, 29°C” condition, estrogen was not administered during starvation (Day 8-10). *n* indicates the number of midgut.

(D) *pros^ts^>T-trace* initially marks Pros^+^esg^−^ cells (arrows).

(E) Quantification of the Pros^+^esg^−^ ratio and Pros^−^esg^+^ ratio in *pros*-lineage cells in T- trace midgut. n=9 (Day 2), 14 (Day 6) midguts.

(F and G) Anterior EEs express more Pgi:GFP protein than do posterior EEs, which is quantified in (G). Day 1 midguts were analyzed. n=97 (Anterior), 109 (Posterior) *pros>mCherry^+^* cells.

(H-L) Candidate screening. *Stat92E, dome, Notch, Tor, Rheb, yki, arm, pan, hep, EGFR,* and *ras85D* were tested. Knockdown of *Stat92E, dome,* and *Notch* significantly increased the number of anti-Pros^+^ cells at Day 3. *n* indicates the number of midguts.

(M) Representative images of *upd3-Gal4>GFP^+^* whole midguts at Day 0 and Day 4 (fed).

(N) Quantification of (M). n=6 (Day 0), 8 (Day 4, fed) anterior midguts.

(O) *upd3-Gal4>GFP* signal is high in non-Pros^+^ cells.

(P) The total cell number increases in the anterior midgut after refeeding. n=15 (Day 10), 14 (Day 11), 14 (Day 12), and 12 (Day 13) guts. The experimental scheme indicated in Figure 7A was applied.

(Q) Representative images of wildtype midgut before/after refeeding. Anti-Pros staining was performed to count the number of Pros^+^ cells (related to Figure 7C).

N.S., not significant: P>0.05, *P≤0.05, **P≤0.01, ***P≤0.001. Two tailed *t* tests (B, E, G, N) and one-way ANOVAs with post hoc Tukey tests (H-L). Scale bars: 20 µm (A), 500 µm (C, M, Q), 50 µm (D, O), 10 µm (F).

