## Supplementary information for "Nutrient-driven dedifferentiation of enteroendocrine cells promotes adaptive intestinal growth"

### Figure S1

Nagai et al.

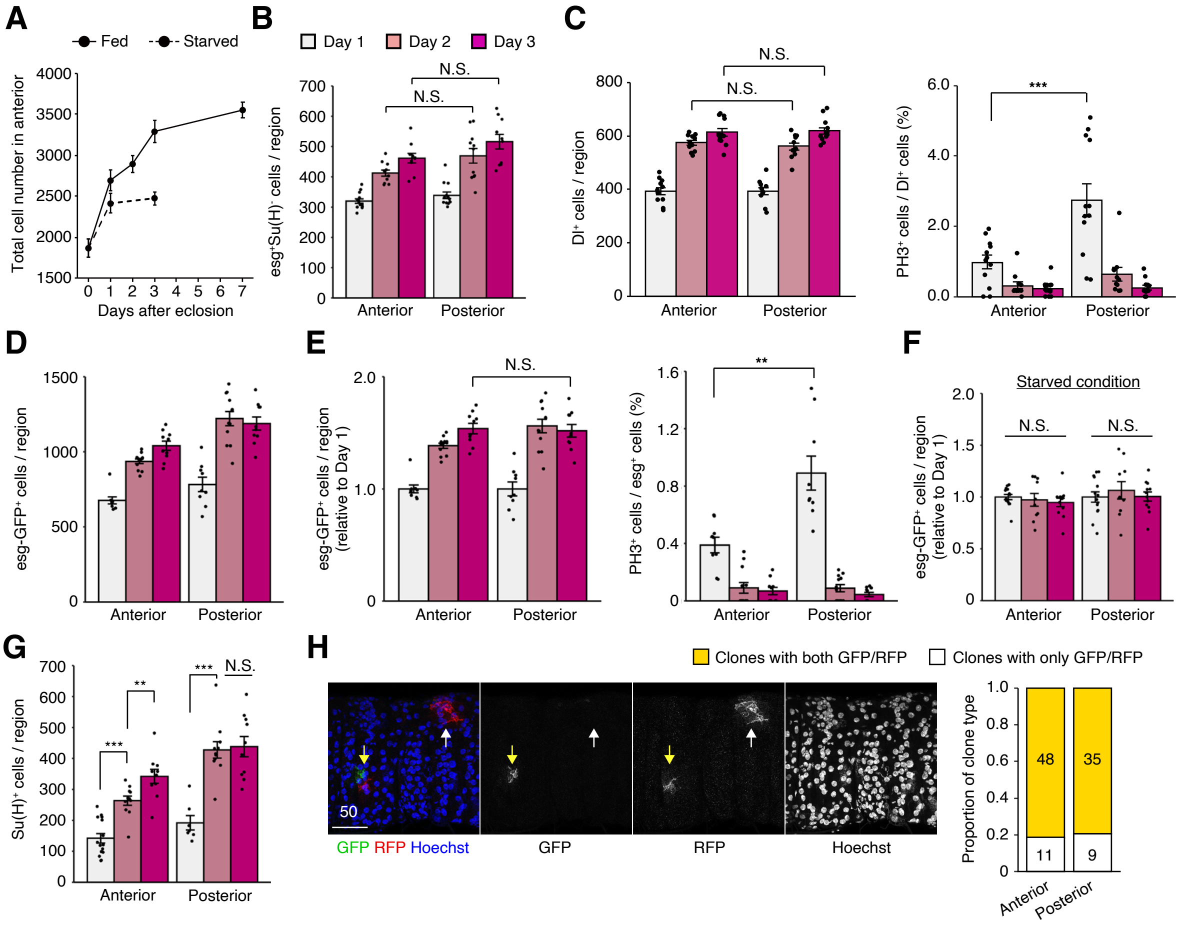

#### Figure S2

Nagai et al.

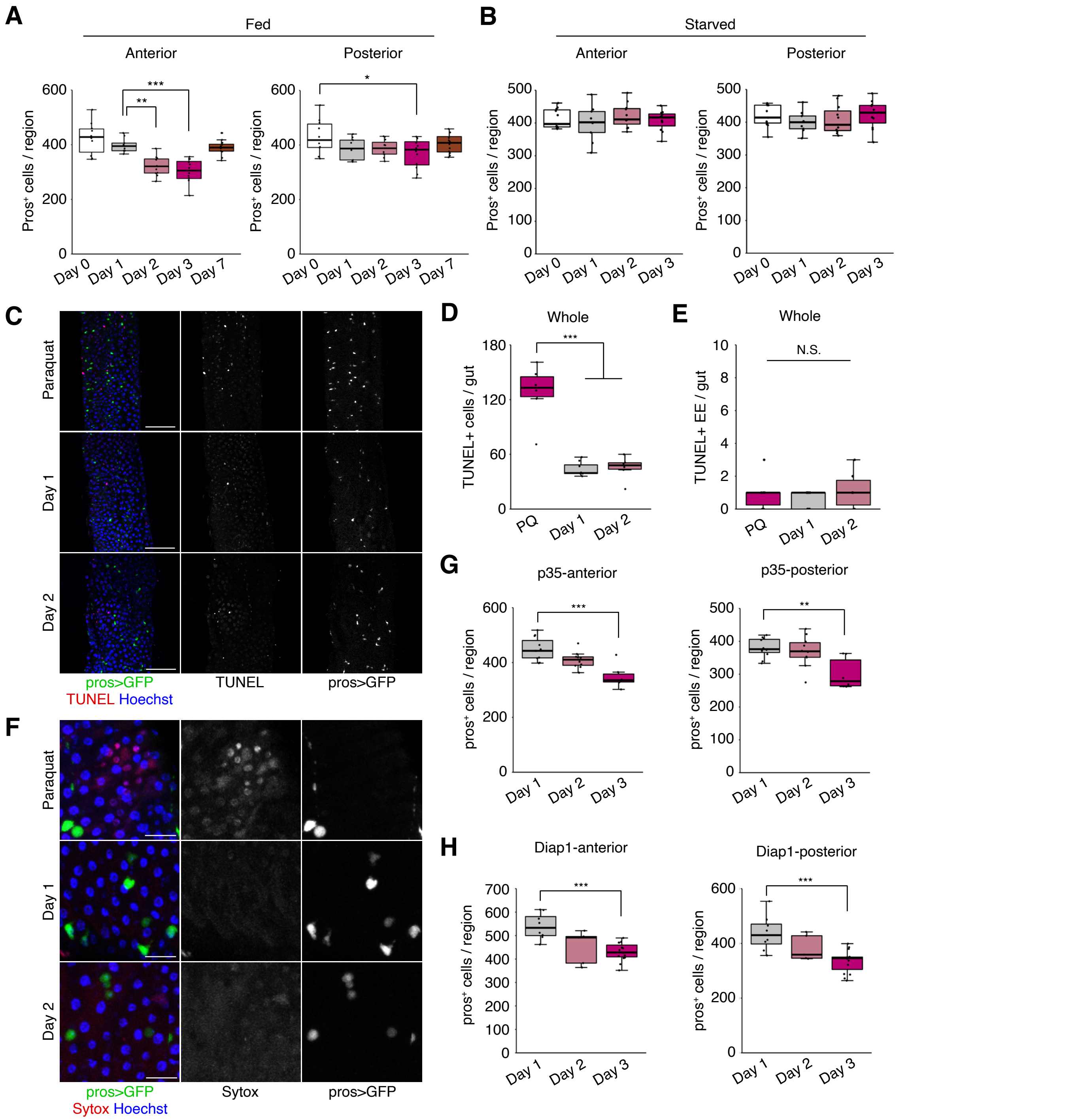

### Figure S3

Nagai et al.

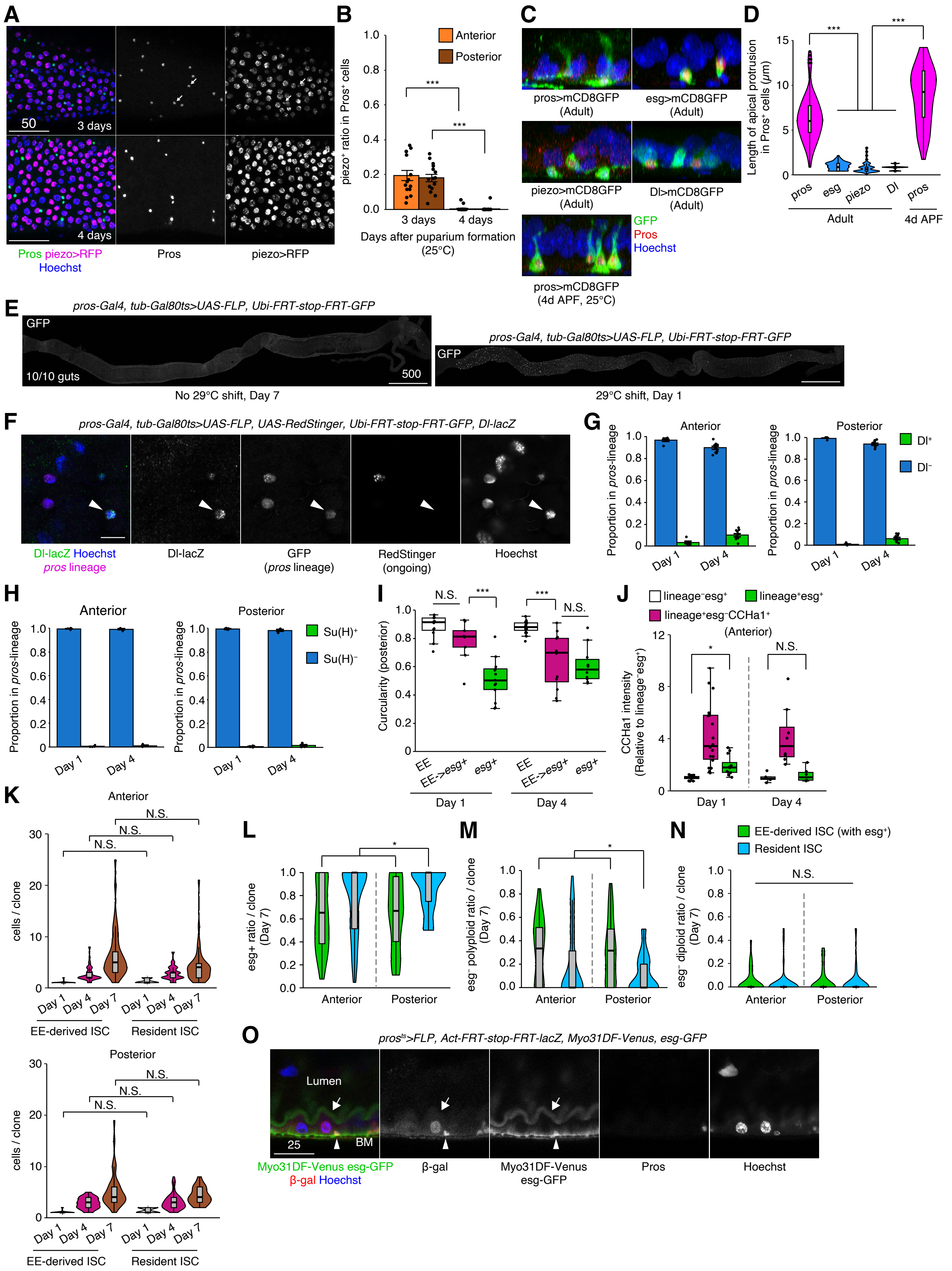

### Figure S4

Nagai et al.

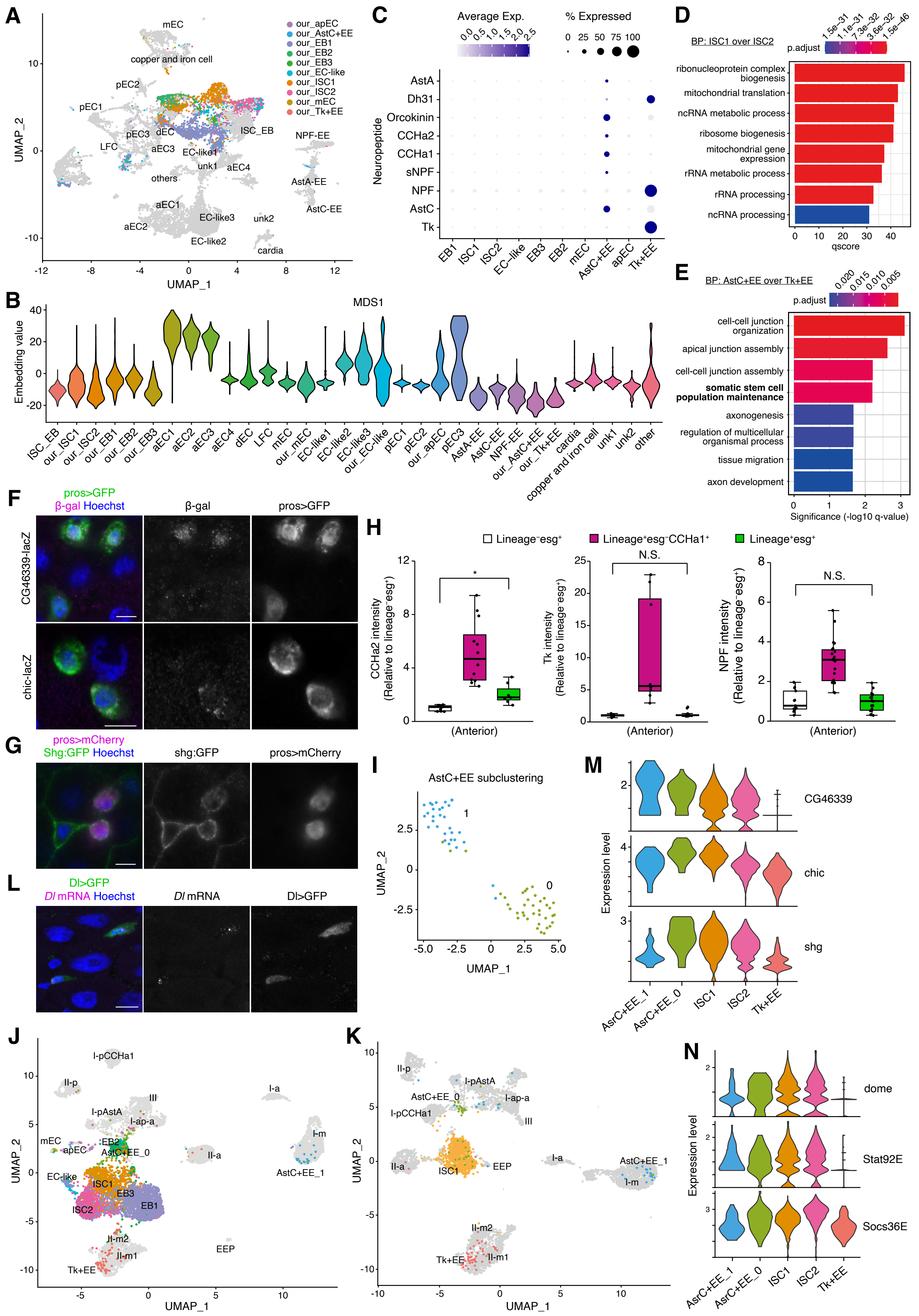

**Figure S5** **Nagai et al.**

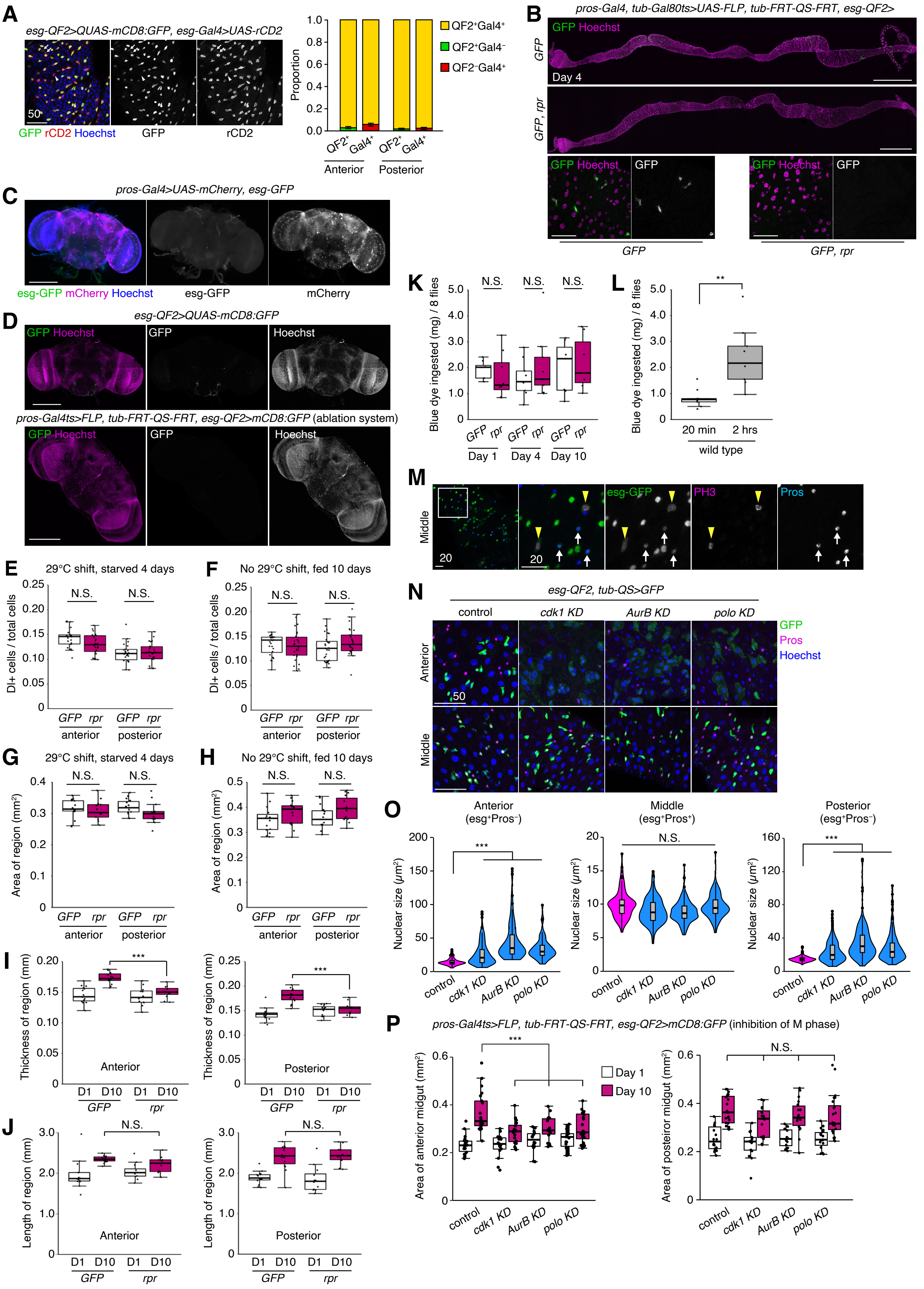

Figure S6 Nagai et al.

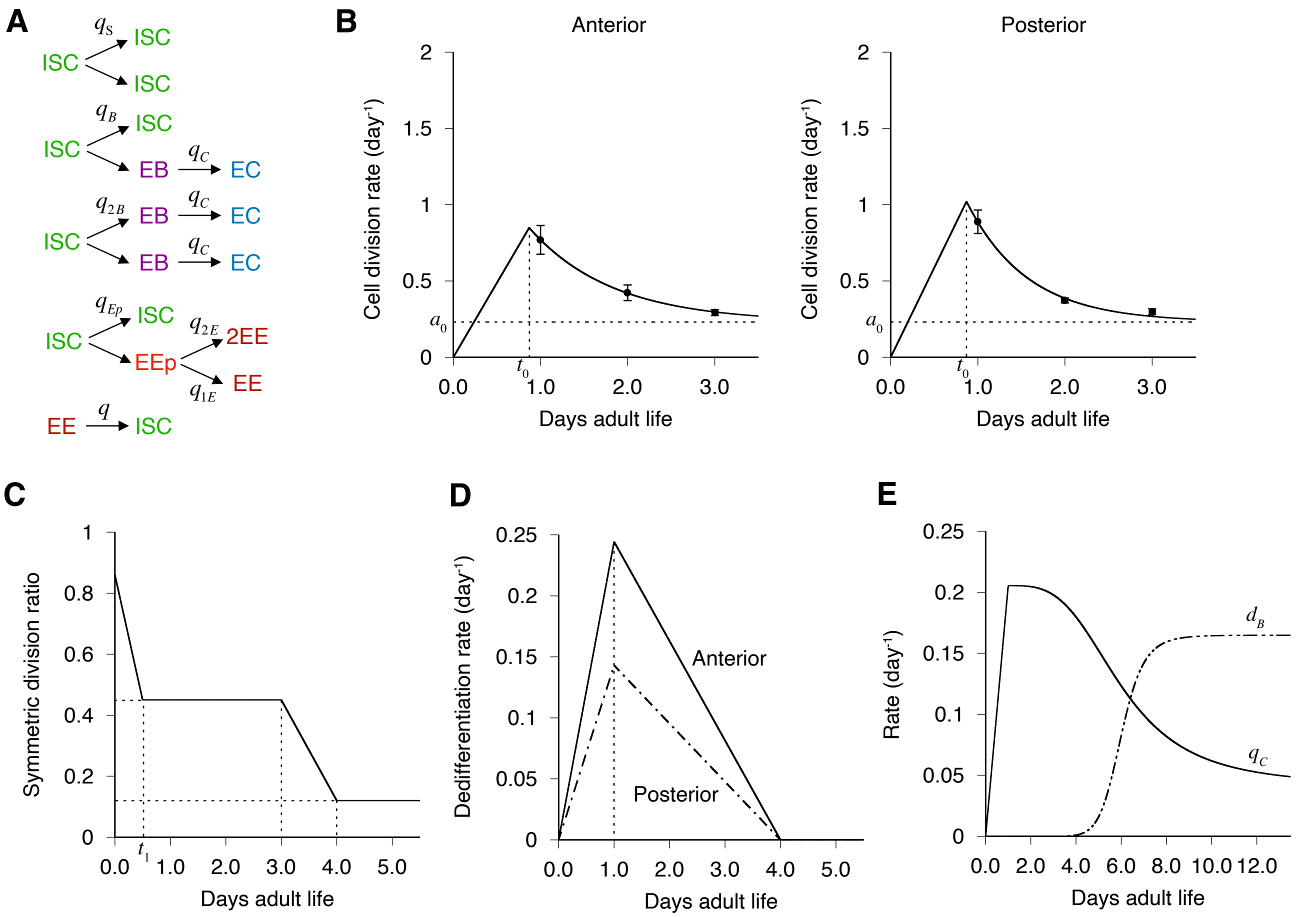

#### Figure S7

Nagai et al.

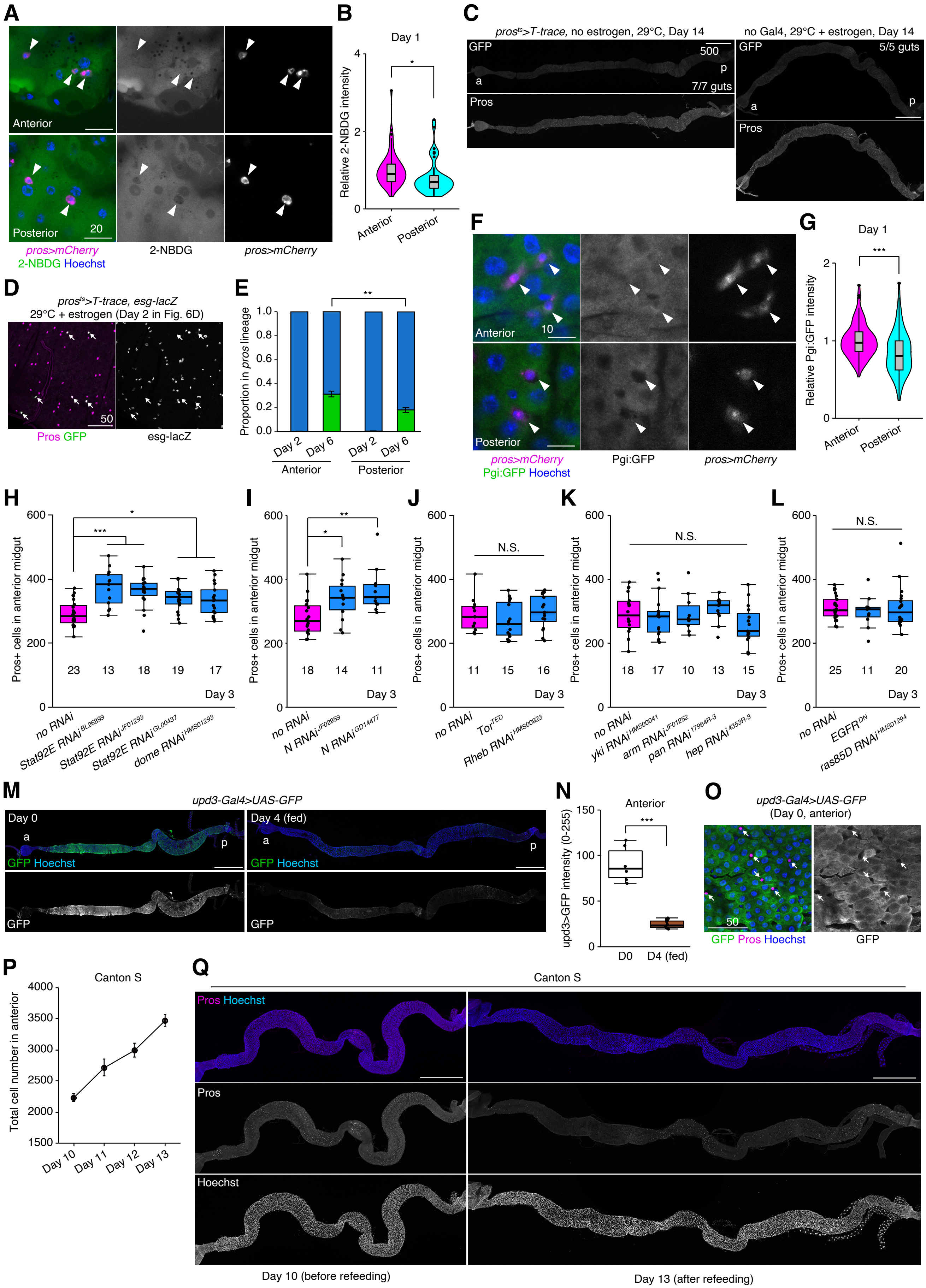
